# Quantitative Modelling of Amyloid-β Dynamics in Brain, CSF, and Plasma During Sleep and Wakefulness

**DOI:** 10.64898/2026.09.07.749991

**Authors:** Satyam Sangeet, Craig L Phillips, Camilla M Hoyos, Angela L D’Rozario, Brendan P Lucey, Svetlana Postnova

## Abstract

**Summary:** Amyloid-beta (Aβ) accumulation in the brain is linked to Alzheimer’s disease. In healthy individuals, Aβ rises during wakefulness and is cleared during sleep, yet the effects of sleep disturbances on the Aβ dynamics are unclear. We developed a model, incorporating brain, cerebrospinal fluid (CSF), and plasma, to investigate the Aβ dynamics along the sleep-wake cycles. The model reproduces the experimentally observed 24-hour Aβ42 oscillations in CSF (660–760 pgml^-1^) and plasma (15–19 pgml^-1^) and predicts brain Aβ dynamics. Indwelling lumbar catheter data show elevated CSF Aβ42 levels on the second morning after a full night of sleep compared to first morning. The model predicts this increased level is due to repeated CSF sampling, which affects Aβ levels through pressure-mediated changes in CSF, impaired sleep-associated clearance, or their synergistic effect. These findings provide a framework for understanding Aβ regulation, highlighting sleep’s protective role and the need for non-invasive measurement approaches.

**Highlights:**

- A quantitative three-compartment model captures 24-hour amyloid-beta dynamics across brain, CSF, and plasma during sleep-wake cycles in healthy individuals.
- The model reproduces normal sleep-wake activity of amyloid-beta in CSF and plasma, consistent with healthy physiological dynamics.
- Model predicts brain amyloid-beta concentrations, providing insights into dynamics experimentally challenging to measure.
- Experimental datasets with repeated lumbar punctures show day-to-day CSF amyloid beta accumulation, deviating from stable dynamics expected under healthy homeostasis.
- The model predicts lumbar punctures alter CSF amyloid beta levels through pressure-mediated changes in CSF, sleep-dependent glymphatic clearance, or both.

## Introduction

Sleep is a fundamental biological process that supports both brain health and cognitive function1,2. Disruptions to normal sleep patterns are increasingly recognised as a major risk factor for neurodegenerative disorders, such as Alzheimer’s disease, where early sleep disturbances often precede cognitive symptoms by years3. This could be explained by the connection between poor sleep and build-up of amyloid beta (Aβ) in the brain4,5. Understanding how sleep disturbances influence Aβ dynamics is critical for exploring the early mechanisms of disease progression and the association between sleep, amyloid dynamics, and cognitive decline. Research has shown that poor sleep both increases Aβ production and impairs its clearance, creating a bidirectional relationship that accelerates accumulation and cognitive decline6. Studying these effects directly in humans is difficult due to the invasive nature of the experimental investigations7. Mathematical modelling provides us with a non-invasive approach to consolidate existing findings and incorporate the influence of sleep-wake cycles on Aβ regulation. Modelling enables us to distinguish between brain interstitial Aβ, CSF Aβ, and plasma Aβ, which experimental methods conflate due to the interconnected nature of these compartments and measurement artifacts introduced by invasive sampling procedures8,9.

In healthy individuals, the central nervous system continuously produces and removes Aβ to maintain protein balance10,11. Neuronal activity regulates Aβ production, with increased cortical activity leading to more Aβ being released into the brain’s interstitial fluid12. The regulation of Aβ levels is a result of several clearance mechanisms, including the blood-cerebrospinal fluid barrier, glymphatic drainage, blood-brain barrier (BBB), and intramural periarterial drainage which work together to regulate Aβ levels13. Also, enzymes like neprilysin and insulin degrading enzyme, along with microglial phagocytosis, help eliminate Aβ locally within the brain parenchyma14,15. Aβ is exchanged between cerebrospinal fluid (CSF) and the bloodstream at the blood-CSF barrier16-19. This exchange is performed by transport proteins with low-density lipoprotein receptor-related protein 1 (LRP1) promoting the clearance of Aβ from the CSF to blood20 and the receptor for advanced glycation end products (RAGE) transporting Aβ in the opposite direction (from blood to CSF)21. The glymphatic system utilises CSF flow along perivascular spaces, driven by arterial pulsations and aquaporin-4 channels on astrocytic endfeet to transport Aβ from interstitial fluid to CSF22-25. Cerebral blood flow dynamics further affect clearance, where impaired perfusion can disrupt Aβ elimination26. Cleared Aβ reaches plasma, where peripheral organs such as the liver and kidneys degrade it, thereby connecting central homeostasis to systemic clearance. Thus, plasma Aβ levels correlate with cerebral amyloid burden27,28, but the strength of this relationship varies by assay platform and the participants involved in the study 29,30.

The sleep-wake cycle has emerged as a critical modulator of Aβ production and clearance, with sleep facilitating the removal of Aβ from the brain through specialised mechanisms31. During wakefulness, increased neuronal activity causes Aβ to be released into the interstitial fluid, with cortical firing rates regulating how the amyloid precursor protein is processed and how Aβ is secreted12. During sleep, CSF and interstitial fluid move through the glymphatic system, which facilitates the removal of neuronal waste products and metabolites32. Studies have reported that increased glymphatic system activity during sleep, particularly slow wave sleep (SWS), which facilitates convective flow of CSF through brain interstitial spaces, helps in the removal of metabolic waste, including Aβ31,33. During SWS, the volume of the interstitial space can increase up to 60%, which reduces the resistance to CSF flow and thereby enhances glymphatic clearance31. Also, neuronal activity and norepinephrine release decrease during SWS, leading to vascular relaxation and increasing interstitial fluid movement, which further supports waste clearance34-36. An alternative hypothesis was recently presented by Miao et al.,37 who used tracer retention measurements and diffusion modelling to demonstrate that brain clearance may be reduced during sleep as opposed to being enhanced. The exact mechanisms thus remain a subject of debate in the field38,39. However, both hypotheses acknowledge that the coordinated cellular and vascular changes along the sleep-wake cycles facilitate physiological clearance of Aβ, therefore reducing the amyloid accumulation33.

Modelling has advanced our understanding of Aβ dynamics by capturing the critical aspects of its production, clearance, aggregation, and transport across brain compartments. Kinetic models have explored Aβ peptide production and exchange within CSF, predicting how rate variations influence concentrations and labelling patterns of Aβ peptides40. Other approaches have treated the brain as a porous medium to describe Aβ polymerisation, highlighting the interstitial fluid’s role in modulating plaque burden41. Stochastic frameworks have simulated aggregation kinetics of Aβ42, while deterministic models have examined the spatial distribution of amyloid deposits and disease progression43,44. At larger scales, network transport models have integrated tau protein dynamics45 and compartmental models have simulated multi-compartmental transport of Aβ46. These models have deepened our understanding of Aβ kinetics, aggregation, and transport. Most recently, Dagum and group developed a six-compartment pharmacokinetic model incorporating sleep-active glymphatic exchange and synaptic-metabolic release and validated it against a randomised crossover clinical trial data measuring overnight changes in plasma Aβ and tau47. The model employs literature-derived kinetic constants to generate the predictions about how sleep versus sleep deprivation alters morning plasma biomarker levels. However, a quantitative framework that simulates the dynamic rather than static effects of sleep and wakefulness on Aβ remains absent. Developing such a model is essential for understanding how cumulative sleep disturbances translate into measurable Aβ burden changes from hours to days.

In this study, we develop a three-compartment model involving the transport between the brain, CSF, and plasma to explore Aβ dynamics under healthy physiological conditions along the sleep-wake cycles. Our model introduces a three-compartmental framework that, unlike prior compartmental approaches46, explicitly incorporates sleep-wake cycle modulation of production and clearance dynamics. We calibrate the model against the experimental Aβ temporal dynamics in CSF and plasma. The calibrated model parameters are then utilised to understand the physiological mechanism affecting the CSF Aβ levels, due to the effect of lumbar sampling.

## Results

### Mathematical Formulation of Sleep-Wake Amyloid-β Dynamics Model

Figure 1 depicts the schematic of our model comprising three interconnected compartments: the Brain, CSF, and Plasma. The model is governed by a set of ordinary differential equations that account for Aβ production and clearance during sleep and wakefulness. In this minimal formulation of the model, we focused on sleep-wake cycles and made the following simplifications: (i) the net flow comprised of forward and back propagation of Aβ between the brain and CSF is modelled with a single net rate term, and (ii) the effect of the circadian drive is present only in the form of the sleep-wake cycles driving Aβ dynamics, so no direct circadian modulation on Aβ metabolism is implemented. The primary objective was to establish a minimal yet mechanistically interpretable model capable of reproducing the established Aβ dynamics.

**Figure 1.**
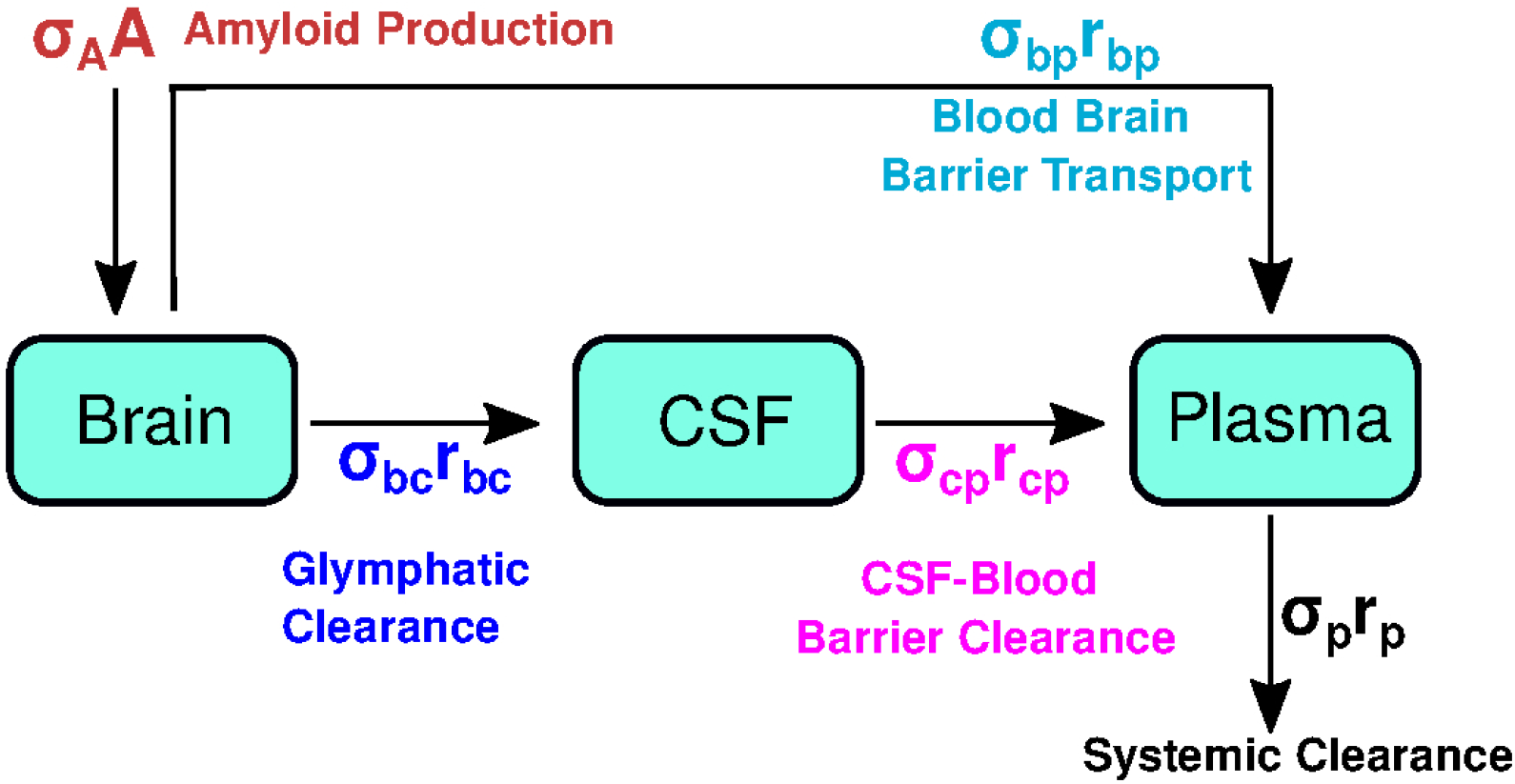
Schematic Representation of the Model. Amyloid is produced in the brain at a rate A (pgml^-1^h^-1^). From the brain, the amyloid is cleared to the CSF at a rate r_bc_ (h^-1^) and to plasma with a rate r_bp_ (h^-1^) during wake. From the CSF, the amyloid is cleared to the plasma at a rate r_cp_ (h^-1^), where the amyloid is removed from the body at a rate r_p_(h^-1^) during wake. All the rates are modulated by the states of sleep and wake through a parameter σ_i_, where i A,bc,bp,cp,p. We assume no backpropagation of amyloid between the compartments and no degradation or other removal pathways except for that from plasma. Circadian control is not explicitly considered in this simplified model; instead, it is accounted for by the state changes between sleep and wake.

#### Brain Compartment

The brain compartment is the primary site of Aβ production and the initial site for the clearance process10,11. Within the brain compartment, the temporal concentration dynamics of Aβ production, denoted as *ρ*_*b*_(*t*) in picograms per millilitre (pgml^-1^), is described as:

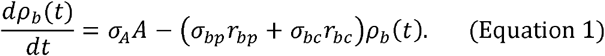

Parameter A represents the amyloid production rate, in pgml^-1^h^-1^. The term *σ*_*A*_*A* captures sleep-wake dependence of amyloid production, with *σ*_*A*_ set to 1 during wake and adjusted during sleep to account for changes in Aβ synthesis50. Clearance from the brain occurs through two distinct pathways: transfer to CSF via glymphatic system represented by *r*_*bc*_ and transfer to plasma via the BBB, represented by *r*_*bp*_. Both these baseline rates are maintained at constant values and are modulated by the sleep-wake state-dependent parameters *σ*31. The effective clearance from the brain compartment (*σ*_*be*_ *r*_*bc*_ + *σ*_*bp*_ *r*_*bp*_) thus adjusts the total clearance dynamically, aligning with the sleep-wake cycle. The assumption of linear first-order kinetics is a useful approximation and is consistent with SILK-based compartmental models used to quantify Aβ production and clearance in humans40,51. However, we acknowledge that Aβ transport across the BBB can exhibit non-linear behaviour, which can be considered in future models, provided more data becomes available 52.

#### CSF Compartment

The CSF compartment acts as an intermediate part in the Aβ clearance pathway. It receives Aβ from the brain and facilitates its transfer to the plasma, with CSF concentration denoted as *ρ*_*c*_ in pgml^-1^. The temporal dynamics of Aβ in CSF are modelled as follows:

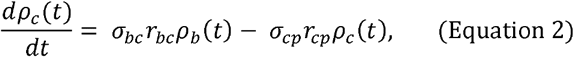

where *σ*_*be*_ *r*_*be*_ *ρ*_*b*_(*t*) denote the influx of Aβ from the brain and *σ*_*cp*_ *r*_*cp*_ *ρ*_*c*_(*t*) models the efflux of Aβ from CSF to plasma, with *r*_*cp*_ as a fixed rate representing the movement across the blood-CSF barrier facilitated by membrane transporters such as LRP119. The sleep-wake dependent factor *σ*_*cp*_ adjusts this rate according to the sleep or wake state. The inclusion of CSF as a distinct compartment, in addition to the direct amyloid transport via BBB, is justified by its role in the glymphatic clearance: Aβ being transported from the brain to CSF and subsequently absorbed in the blood33,53.

#### Plasma Compartment

The plasma compartment represents the systemic circulation, where Aβ is finally removed from the body. The dynamics of Aβ concentration in the plasma compartment, ρ_*P*_ are governed by:

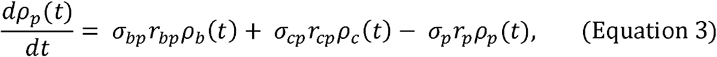

where the influx terms, *σ*_*bp*_ *r*_*bp*_ *ρ*_*b*_(*t*) and *σ*_*cp*_ *r*_*cp*_ *ρ*_*c*_(*t*) represent the movement of Aβ from the brain to plasma via the BBB and CSF to plasma via the blood-CSF barrier, respectively. The efflux term, *σ*_*p*_ *r*_*p*_ *ρ*_*p*_(*t*) models the removal of Aβ from plasma via renal excretion54, hepatic metabolism55, and peripheral tissue absorption56. The *σ*_*p*_ factor dynamically adjusts this clearance rate in response to sleep-wake cycles, reflecting physiological differences in systemic processing. While direct experimental evidence for sleep-dependent modulation of systemic plasma clearance is currently lacking, sleep dependent variation in hepatic and renal enzyme activity provides biological plausibility for the sleep dependence of systemic clearance57-59. The inclusion of the plasma compartment is physiologically justified by its role as the final stage of Aβ elimination20, with plasma Aβ levels correlating with cerebral amyloid burden60.

#### Model Steady States and Solutions

The model presented in Equations 1-3 is a piecewise linear model, which means that in each phase, wake or sleep, the solutions for *ρ*_*b*_(*t*), *ρ*_*c*_(*t*) and *ρ*_*p*_(*t*) tend towards that phase’s steady state. The steady-state concentrations during wake and sleep are:

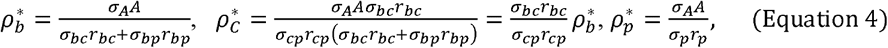

where all *σ* are equal to 1 during wake. The amyloid production rate *σ*_*A*_*A* scales all steady-state concentrations. The steady-state concentration in the brain is affected only by *A, r*_*bc*_,and *r*_*bp*_, and their scaling factors. The CSF steady state is proportional to that in the brain via 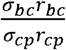. This implies that if the clearance from the brain to CSF is faster than from CSF to plasma, then the CSF steady-state concentration is higher than in the brain and vice versa. The plasma steady state is affected only by *A* and *r*_*p*_ and their scaling factors.

These steady-state concentrations define the boundaries within which Aβ concentrations fluctuate during the sleep-wake cycles. When switching between sleep and wake, the model switches to a new target steady state. If a wake steady state concentration is higher than the sleep one, then Aβ concentration in the given compartment increases during wake and decreases during sleep. The opposite is true if the sleep steady-state concentration is higher than the wake one. For example, in plasma, this can be achieved if 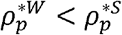, which is possible only if 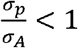. For completeness, results of the model sensitivity analysis and the effects of varying parameters are shown in Figures S1-S3.

The time-dependent solutions in each state are then:

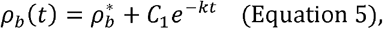

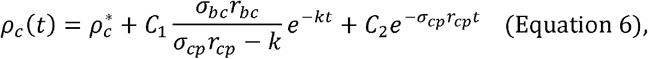

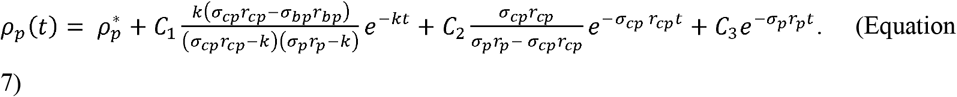

Where *k* = *σ*_*bp*_ *r*_*bp*_ + *σ*_*bc*_ *r*_*bc*_, and all are equal to 1 during wake. The constants *C*_1_,*C*_2_, and *C*_3_ are determined by the initial conditions for the concentrations and define how far the system starts from the steady state. The brain concentration follows an exponential rise/decay depending on whether the value is above or below the steady state. The concentrations in CSF and plasma follow a multi-exponential rise/decay. The exponents in the solutions above are always negative because the transfer rates and scaling factors are physiologically constrained to be positive. As a result, the model is stable for any regular sleep-wake cycle. Regardless of the initial conditions, it will eventually lock into a stable cycle following the repeating sleep-wake times.

The intrinsic time constants for each compartment are:

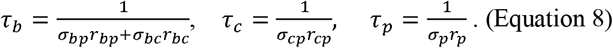

Due to the cascade structure of the model, the time it takes for *ρ*_*p*_ (*t*) to stabilise is governed purely by *τ*_*b*_, the time it takes for *ρ*_*c*_ (*t*) depends on both *τ* _*b*_ and *τ*_*c*_, and the time it takes for *ρ*_*p*_ (*t*) to stabilise depends on all three intrinsic time constants.

### Parameter Constraints

To ensure that an optimal parameter set can be identified by fitting the model to the data, we first need to constrain the parameter ranges within which they may vary. Specific boundaries and conditions based on experimental data were applied to model parameters where available. We first fit the model to Aβ42 dynamics and then adjust it for Aβ40. The resulting parameter constraints and the fitted values for Aβ42 are provided in Table 1.

**Table 1.** Parameter values and Constraints for A*β*42 under Normal Sleep-wake Cycles.

| Parameter | Default | Blattner et al 2020 | Lucey et al 2018 | Liu et al 2023 | Meaning, units | Experimental data | Constraint based on data | Fitting bounds |
| --- | --- | --- | --- | --- | --- | --- | --- | --- |
| **Production rate during wake** |  |  |  |  |  |  |  |  |
| $A$ | 16.203 | 84.523 | 55.780 | 14.450 | $A\beta$ production in the brain, [pgml<sup>-1</sup>h<sup>-1</sup>] | - | $A = \frac{r_p(T_w \rho_p^w + T_s \sigma_p \rho_p^s)}{T_w + T_s \sigma_A} \leq 111$<br>[pgml<sup>-1</sup>h<sup>-1</sup>] | 0-111 |
| **Clearance rates during wake** |  |  |  |  |  |  |  |  |
| $r_{bc}$ | 0.038 | 0.019 | 0.062 | 0.015 | brain to CSF, [h<sup>-1</sup>]. | - | $r_{bc} + r_{bp} < 0.25 \text{ h}^{-1}$ | 0.01-0.25 |
| $r_{bp}$ | 0.014 | 0.034 | 0.040 | 0.014 | brain to plasma, [h<sup>-1</sup>] | - | $r_{bc} + r_{bp} < 0.25 \text{ h}^{-1}$ | 0.01-0.25 |
| $r_{cp}$ | $0.537 \times 10^{-2}$ | $1.54 \times 10^{-2}$ | $1.56 \times 10^{-2}$ | $0.320 \times 10^{-2}$ | CSF to plasma, [h<sup>-1</sup>] | - | $r_{cp} < \frac{A}{\rho_c^{wake}} < 0.1 \text{ h}^{-1}$ | 0-0.1 |
| $r_p$ | 0.427 | 0.298 | 0.300 | 0.475 | from plasma out of the body, [h<sup>-1</sup>] | Half-life of $A\beta$ in plasma is around 2-3 hours (Ovod et al., 2017), resulting in clearance of 0.23 to 0.34 h<sup>-1</sup> | $r_p = 0.231 - 0.347 \text{ h}^{-1}$ | 0-0.6 |
| **Scaling factors for production and clearance during sleep** |  |  |  |  |  |  |  |  |
| $\sigma_A$ | 0.772 | 0.633 | 0.485 | 0.750 | Scaling production, [-]. | Amyloid production reduces by 10-30% during sleep (DiNuzzo et al., 2021) | $\sigma_A < 1$ | 0-0.99 |
| $\sigma_{bc}$ | 1.131 | 1.660 | 1.002 | 1.110 | brain to CSF, [-]. | 2.17-fold increase in clearance during sleep in mouse (Xie et al., 2013). | | 1-7 |
| $\sigma_{bp}$ | 1.768 | 1.816 | 2.479 | 1.297 | brain to plasma, [-] | - | | 1-7 |
| $\sigma_{cp}$ | 6.100 | 5.740 | 4.087 | 6.552 | CSF to plasma, [-] | - | | 1-7 |
| $\sigma_p$ | 4.253 | 3.610 | 4.000 | 3.055 | From plasma out of the body, [-] | - | | 1-7 |
| **Goodness of Fit** |  |  |  |  |  |  |  |  |
| NRMSE | 0.247 | 0.137 | 0.128 | 0.209 |  |  |  |  |

#### Systemic plasma clearance, r_**p**_

Systemic clearance rate was constrained by the experimentally reported plasma half-life of Aβ. The half-life (*τ*_1/2_) is directly related to the clearance rate and represents the time required for Aβ to decrease to half of its initial value. The half-life of Aβ42 in plasma during wakefulness is reported to be between two and three hours61, providing a constraint for the plasma clearance rate, *r*_*p*_:

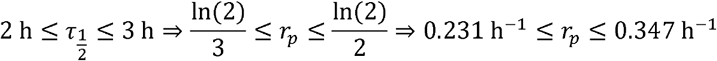

We used the mean value of this range (*r*_*p*_ = 0.285 h^-1^) as the starting parameter value and applied ±100% variation to allow for wider variability, giving the final constrained range for *r*_*p*_ as [0, 0.6] h^-1^.

#### Sleep-dependent scaling of amyloid production, σ_**A**_

Compared to wakefulness, the neuronal firing rate in the cortex drops by 10% to 30% during sleep50. Although these data are from an experiment conducted in mice62, assuming that neural activity produces amyloids, we can propose that amyloid production would be reduced during sleep compared to wake in humans. This introduces a constraint on *σ*_*A*_ < 1; e.g., a value of 0.8 was used in Tekieh et al., 202263.

#### Sleep-dependent scaling of the clearance rates, σ_bc_, σ_bp_, σ_cp_ and σ_p_

The sleep-dependent scaling factors *σ*_*bc*_, *σ*_*bp*_, *σ*_*cp*_ and *σ*_p_ were constrained to the range of [1,7]. As sleep enhances the clearance of Aβ via glymphatic clearance31, across the BBB64, from CSF to plasma24, and systemic circulation65, *σ*_*bc*_, *σ*_*bp*_, *σ*_*cp*_ and *σ*_*p*_ were constrained to be greater than 1. In the absence of more precise experimental data, the upper bound of 7 was chosen to limit the parameter search space to physiologically realistic values. This implies that clearance cannot accelerate more than 7 times during sleep compared to wake. For example, Xie et al.31, reported a 2-fold increase in glymphatic clearance during sleep.

#### Amyloid production rate during wakefulness, A

Amyloid production governs concentrations in all three compartments. Adding all three concentration equations together, Equations 1-3, the internal transfer terms cancel out, giving the total mass balance:

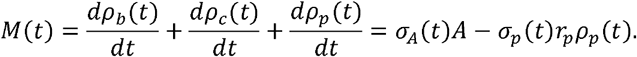

The system is expected to exhibit a stable periodic behaviour over 24 hours, so the amyloid mass that has been produced must be equal to the cleared mass, resulting in a zero net effect:

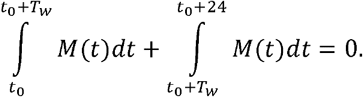

This mass balance assumes that all produced Aβ remains in soluble form and is cleared through clearance pathways and does not account for the clearance of Aβ as insoluble plaques. This assumption is appropriate for the cognitively normal, amyloid-negative individuals, in whom the plaque burden is negligible18. But this limits the model’s applicability to amyloid-positive individuals where plaque deposition represents a sink for Aβ42 deposition. Here *T*_*W*_ is the wake duration, and sleep duration is defined as *T*_*s*_ = 24 − *T*_*W*_. Substituting the value of *M*(*t*) from Equation (8) into Equation (9), using 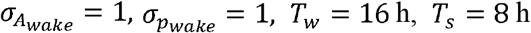, and solving for *A*, we get

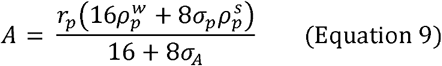

where 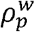 and 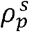 are the concentrations achieved at the end of the wake and sleep states, respectively. To establish an upper bound for *A*, we need to maximise the right-hand side of Equation 8. Substituting the maximum experimentally observed value of amyloid concentration in plasma (41pgml^-1^, see Supplementary Material) and parameter constraints for *r*_*p*_, *σ*_*A*_,and *σ*_*p*_ from Table 1, give us the upper bound for *A* ≤ 111 pgml^-1^h^-1^.

#### Brain to CSF and Brain to Plasma clearance during wakefulness, r_bc_ and r_bp_

*r*_*bc*_ and *r*_*bp*_ represent the net wake state fluxes from brain-to-CSF and brain-to-plasma respectively, which implicitly considers any reverse transport from CSF or plasma into the brain. Direct experimental estimates for *r*_*bp*_, *r*_*bp*_ are unavailable. We constrained these rates by requiring that brain Aβ concentrations do not reach steady state within a single 16-hour wake or 8-hour sleep episodes. For first-order kinetics, reaching 98% of steady-state requires four time constants66. This translates into the requirements for 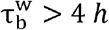 and 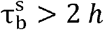. Using Equations 6 and 7, this yields:

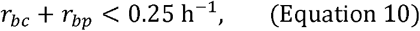

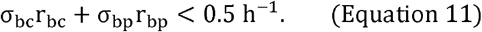

Lower bounds of *r*_*bc*_, *r*_*bp*_ ≥ 2 0.01 h^-1^ were imposed so that the optimisation doesn’t yield unphysiological parameter values. Lower bounds serve as a physiological regularization rather than arbitrary limits. Since these constrains acts on sums rather than individual parameters, we used numerical bounds *r*_*bc*_, *r*_*bp*_ ∈ [0.01,0.25] h^-1^, enforcing constraints (Equations 9-10) during model fitting (see Methods).

#### CSF to Plasma clearance during wakefulness, r_cp_

During wakefulness, the CSF Aβ steady state is 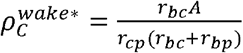, eq 4. Re-arranging for *r*_*cp*_, and accounting for the facts that (i) all clearance rates must be positive, (so 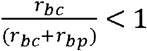), (ii) the maximum observed concentration of Aβ42 in CSF based on experimental data is 1200 pgml^-1^(see Supplementary) and (iii) A ≤ 111 pgml^-1^h^-1^ we get 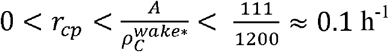 . Thus, *r*_*cp*_ is constrained to [0, 0.1] h^-1^. It is important to note that *r*_*cp*_ represents the net CSF-to-plasma efflux, implicitly capturing the balance between forward efflux and RAGE-mediated re-entry from plasma.

### Normal Sleep-Wake Cycle and Aβ42 Homeostasis

First, we identify the default parameter set to represent the dynamics during normal sleep-wake cycles (16 h diurnal wake and 8 h of nocturnal sleep). The following experimental studies met our inclusion criteria for model fitting: Blattner et al.48 (CSF-only data), Lucey et al.4 (CSF-only data), and Liu et al.49 (simultaneous CSF and plasma). We assume that Aβ42 homeostasis is preserved during normal sleep-wake cycles, meaning that Aβ42 dynamics are stable from one day to the next. Because CSF data are typically affected by the lumbar puncture procedure67,68, preventing normal clearance of Aβ42 during sleep, we detrended the data to reconstruct two subsequent days of stable data from the normal wake dynamics on day 1 (see Methods). The default parameter set was obtained by fitting the model to all available data simultaneously (global fit). We also fitted the model to each study separately to obtain the study-specific parameter sets.

#### Model Dynamics

The default and individual parameter sets resulting in the best fit to normal sleep-wake cycles data are listed in Table 1, and the resulting dynamics for Aβ42 are shown in Figure 2. The default parameter set achieves a global NRMSE of 0.247 across all datasets, with individual fits ranging from 0.128 to 0.209, maintaining the characteristic oscillation with Aβ42 concentrations declining by approximately 13% in CSF and ∼16% in plasma during the sleep phase. Residual analysis on individual fits confirmed mean errors of less than 0.6% of mean compartmental concentrations, with no systematic temporal structure across any dataset (Fig. S4).

**Figure 2.**
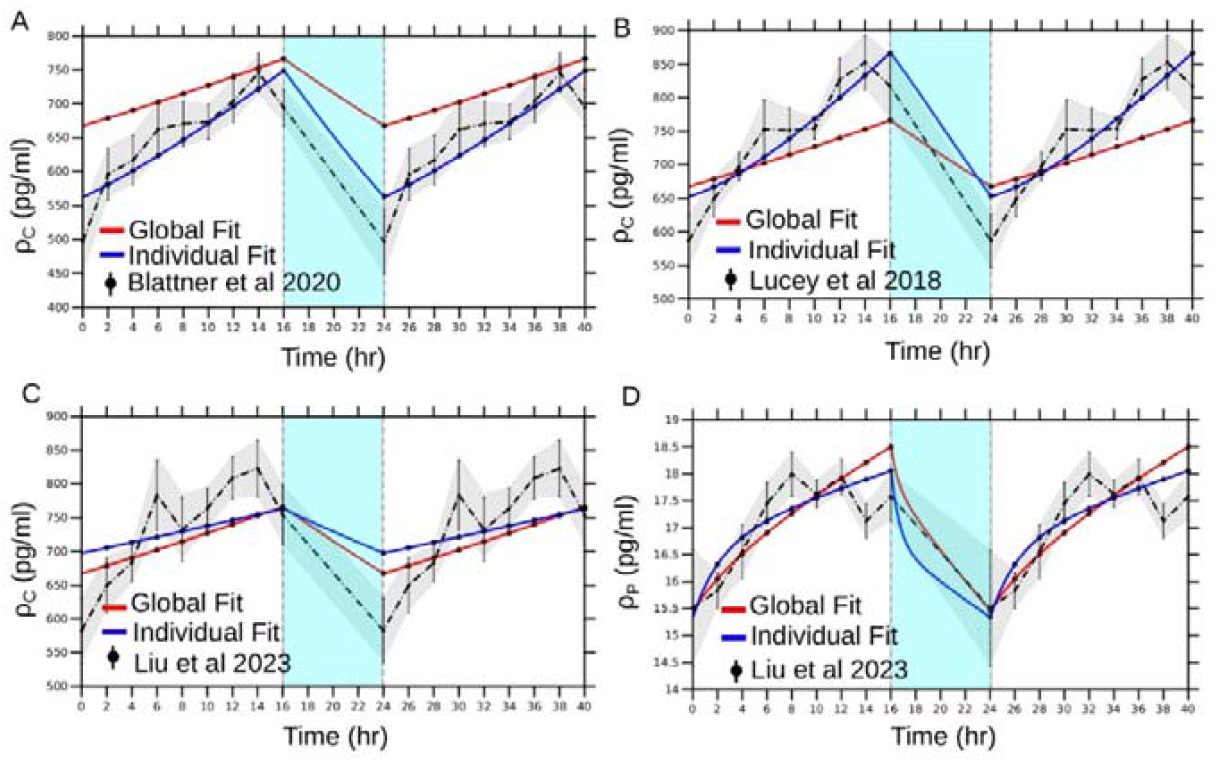
Model Fitting for Normal Sleep-Wake Cycle. **(A)** CSF data (Blattner et al., 2020)^13^ **(B)** CSF data (Lucey et al., 2018)^14^ **(C)** CSF data (Liu et al., 2023)^15^ **(D)** Plasma data (Liu et al., 2023)^15^. The grey shaded area denotes the experimental standard deviation. The cyan shaded area corresponds to the sleep state. The vertical dashed line at t=16 and t=24 denotes the onset and ending of sleep state. For Liu et al., 2023 dataset (containing both CSF and Plasma Aβ42 reading), the model fitting was performed simultaneously to both datasets.

Fitting the model for the individual studies improves the goodness of fit, with NRMSE decreasing for all three studies. For the CSF-only datasets (Blattner et al.48 and Lucey et al.4), the individual fits reproduced the sharper initial rise in CSF Aβ42 concentrations during wakefulness in Figure 2a, 2b, which was not captured during the default fit. This improvement was mainly driven by increased Aβ42 production, *A*, and increased CSF-to-plasma wake state clearance, *r*_*cp*_, in both datasets, Table 1. For Blattner et al.48, the improved fit was mainly driven by an increase in production rate (*A*) and higher sleep-phase brain-to-CSF clearance, capturing the sharper initial CSF Aβ42 rise during wakefulness (Table 1). For Lucey et al.4, similar parameter changes were observed (Table 1).

Conversely, for Liu et al.49 (Figure 2c, 2d), the individual fits resulted in the reduction of *A* from 16.2 pgml^-1^h^-1^ to 14.4 pgml^-1^h^-1^ likely because of the simultaneous fitting of CSF and plasma. Wake phase clearance rates *r*_*bc*_, *r*_*cp*_ decreased and *r*_*p*_ increased compared to the default (Table 1), while sleep phase clearance scaling factors ( *σ*_*bc*_, *σ*_*cp*_) remained relatively similar to the default parameter values, with *σ*_*bp*_, *σ*_*p*_ showing a reduction. This lower wake-state clearance and similar sleep scaling parameters maintain the relative oscillation amplitude.

The model dynamics across all three compartments using the default parameter set are shown in Figure 3. The model reproduces the sleep-wake oscillations of Aβ42 in CSF and plasma compartments in agreement with experimental concentration ranges, while predicting the Aβ42 dynamics in the brain compartment. In the brain compartment in Figure 3a, Aβ42 levels rise during the wake state and decline during sleep, demonstrating an oscillatory pattern between 240 and 280 pgml^−l^. The CSF compartment in Figure 3b exhibits a similar pattern but with larger amplitude fluctuation between 667-770 pgml^−l^. Plasma compartment in Figure 3c mirrors the CSF dynamics but with smaller absolute fluctuations, 15-19 pgml^−l^.

**Figure 3.**
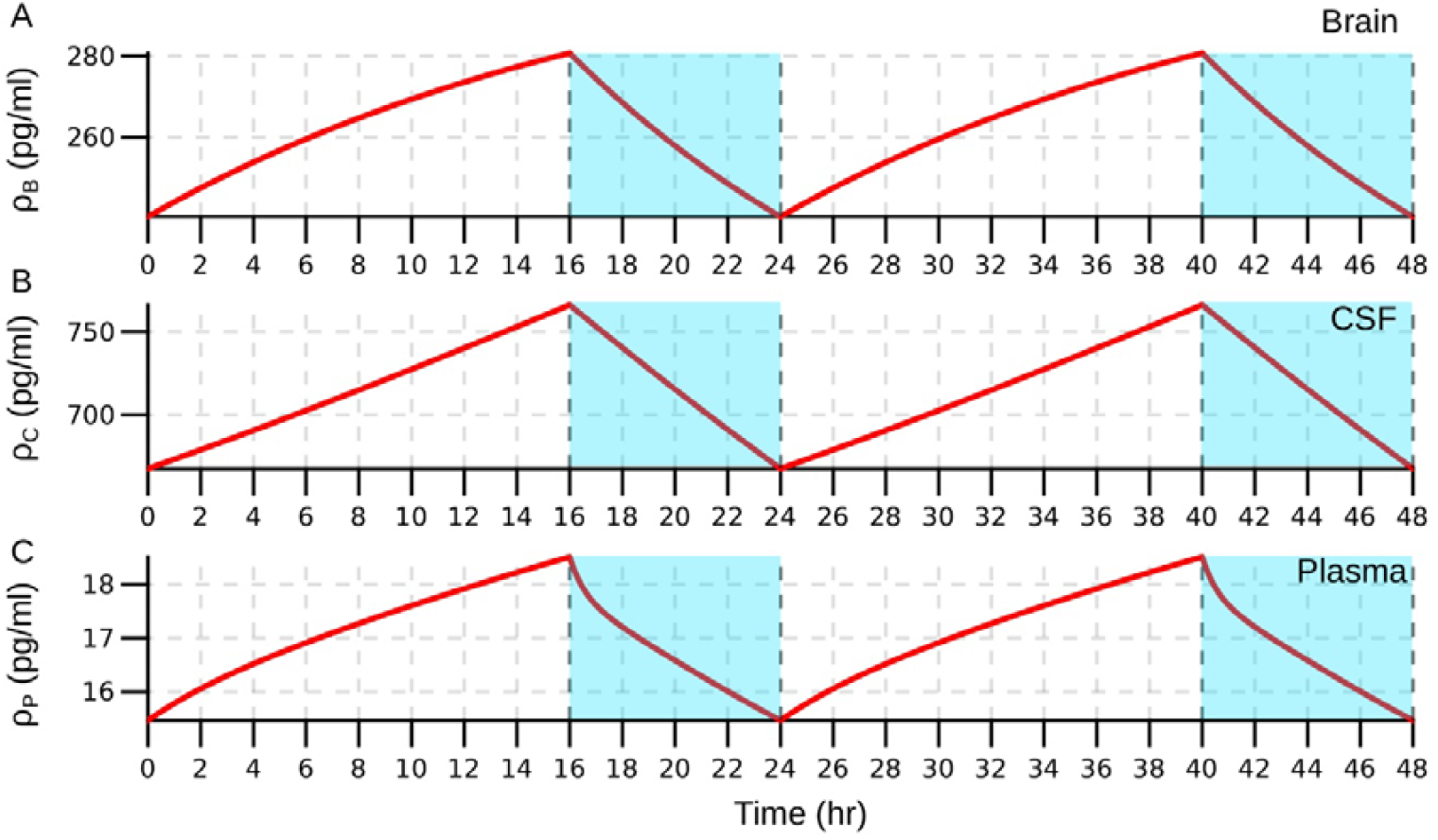
Model Aβ42 Dynamics over a Period of 48 Hours. **(A)** The brain amyloid concentration fluctuates between 225-285 pg/ml. **(B)** The CSF amyloid concentration fluctuates between 670-790 pg/ml. **(C)** The plasma amyloid concentration fluctuates between 15 – 19 pg/ml. The cyan shaded area corresponds to sleep state. The vertical dashed line corresponds to the wake-to-sleep transition (t=16 hr, t=40 hr) and the sleep-to-wake transition (t=24 hr, t=48 hr)

### Effects of lumbar puncture on Aβ42 clearance in CSF

#### Lumbar Puncture Effect

The 36-hour sleep-wake experiments in Blattner et al.48, Lucey et al.4 and Liu et al.49, report elevated CSF Aβ42 levels after awakening from a full night of sleep on day 2 compared to day 1, as demonstrated in Figure 4 and Figure S5. If observed during the normal sleep-wake cycles, such dynamics would lead to a continuous accumulation of Aβ42, quickly resulting in levels inconsistent with healthy functioning. We propose that these dynamics are primarily caused by the indwelling lumbar catheter procedure interfering with the normal clearance process and are not present in an unperturbed case. We propose three potential mechanisms underlying this CSF Aβ42 increase in the experiments with lumbar puncture:

**Figure 4.**
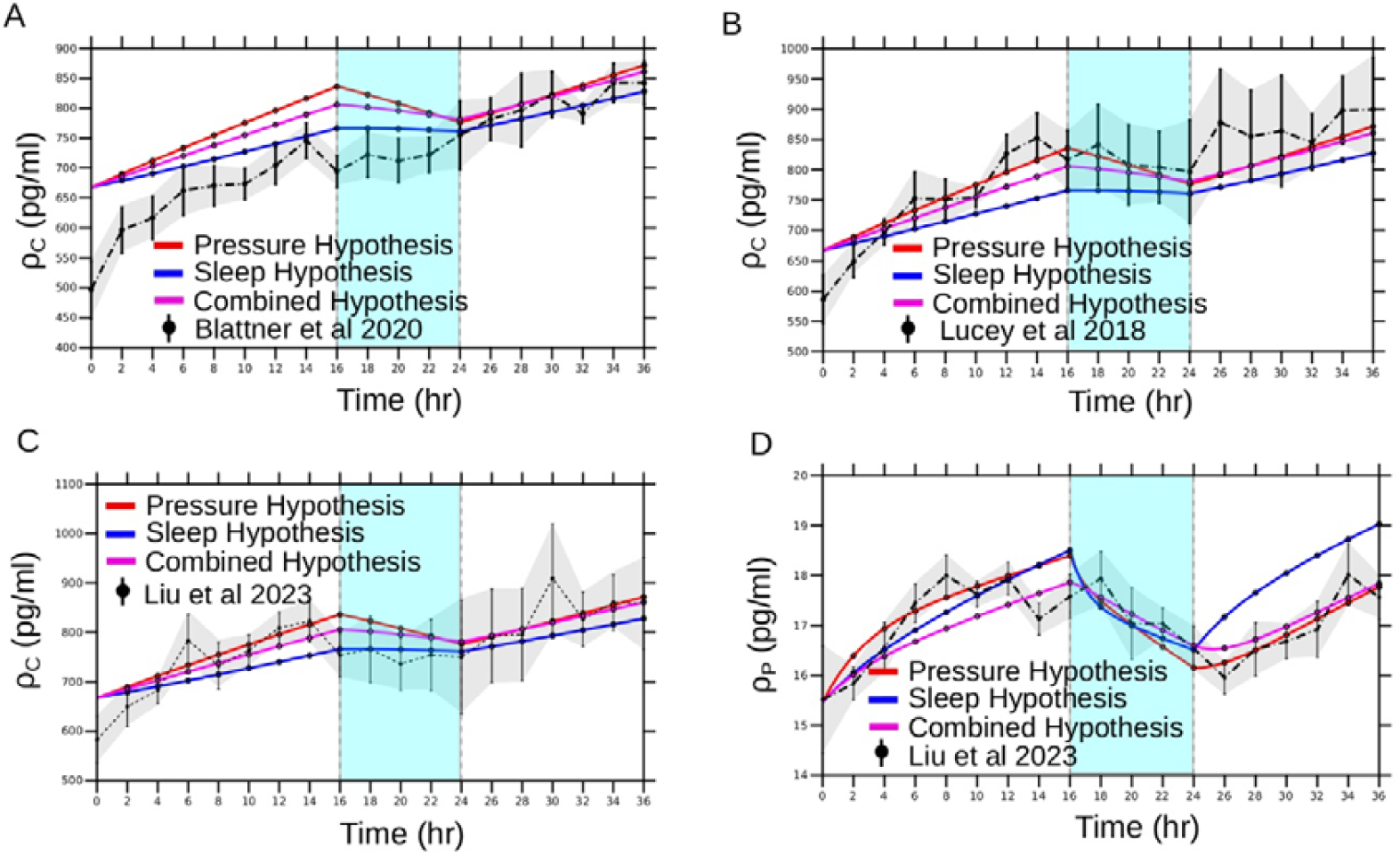
Model Fit to Experimental Data. The red curve corresponds to the best parameter combination fit for the pressure hypothesis (optimising *r*_*bc*_, *r*_*cp*_), the blue curve corresponds to the best parameter combination fit for the sleep hypothesis (optimising *σ*_*bc*_, *σ*_*cp*_ ) and the magenta curve corresponds to the best parameter combination fit for the combined pressure and sleep hypothesis (optimising *r*_*bc*_, *σ*_*bc*_, *σ*_*cp*_ ) **(A)** CSF data (Blattner et al., 2020) **(B)** CSF data (Lucey et al., 2018) **(C)** CSF data (Liu et al., 2023) **(D)** Plasma data (Liu et al., 2023). The red curve corresponds to the pressure gradient hypothesis fit, the blue curve corresponds to sleep disruption hypothesis fit, and the magenta curve corresponds to the combined hypothesis fit. The grey shaded area denotes the experimental standard deviation. The cyan shaded area corresponds to the sleep state. The vertical dashed line at t=16 and t=24 denotes the onset and ending of the sleep state.

1. Pressure hypothesis: The removal of 6ml CSF samples every 2 hours, as done in the experimental studies, is likely to reduce the total CSF volume and thus also reduce the intracranial pressure67,68. This creates a pressure gradient between the brain and CSF compartments, which could increase *r*_*bc*_ (brain-to-CSF clearance), but reduce *r*_*cp*_ (CSF-to-plasma clearance). Reduced intracranial pressure may also affect *r*_*bp*_ (brain-to-plasma clearance) and/or *r*_*p*_ (plasma-to-systemic circulation) if compensatory hemodynamic changes occur69,70. Previous studies have demonstrated that repeated lumbar punctures within 3 days may affect CSF biomarker levels71, supporting the pressure hypothesis. To test these mechanisms, we fitted the model to data using the six most physiologically relevant combinations of *r*_*bc*_,*r*_*cp*_,*r*_*bp*_,*and r*_*p*_, as shown in Table 2, while keeping all other parameters constant.
2. Sleep hypothesis: The invasive nature of lumbar punctures likely disrupts normal sleep architecture and quality, directly impacting the glymphatic clearance. Sleep enhances glymphatic31, BBB64, blood-CSF33, and systemic clearance65 while reducing neural activity50 and thereby suppressing Aβ42 production. Therefore, sleep disruption may impair all these clearance mechanisms through reduced *σ* _*bc*_, *σ* _*bp*_, *σ* _*cp*_, *σ* _*p*_, and increased *σ* _*A*_ . To test which sleep-dependent mechanisms are disrupted by lumbar punctures, we evaluated 18 most relevant parameter combinations as shown in Table 2.
3. Combined Pressure and Sleep hypothesis: Lumbar punctures may simultaneously induce both pressure changes and sleep disruption. To test whether pressure and sleep effects act synergistically, we evaluated the model’s fits to data using seven parameter combinations based on best fits found for each hypothesis separately as shown in Table 2.

**Table 2.**
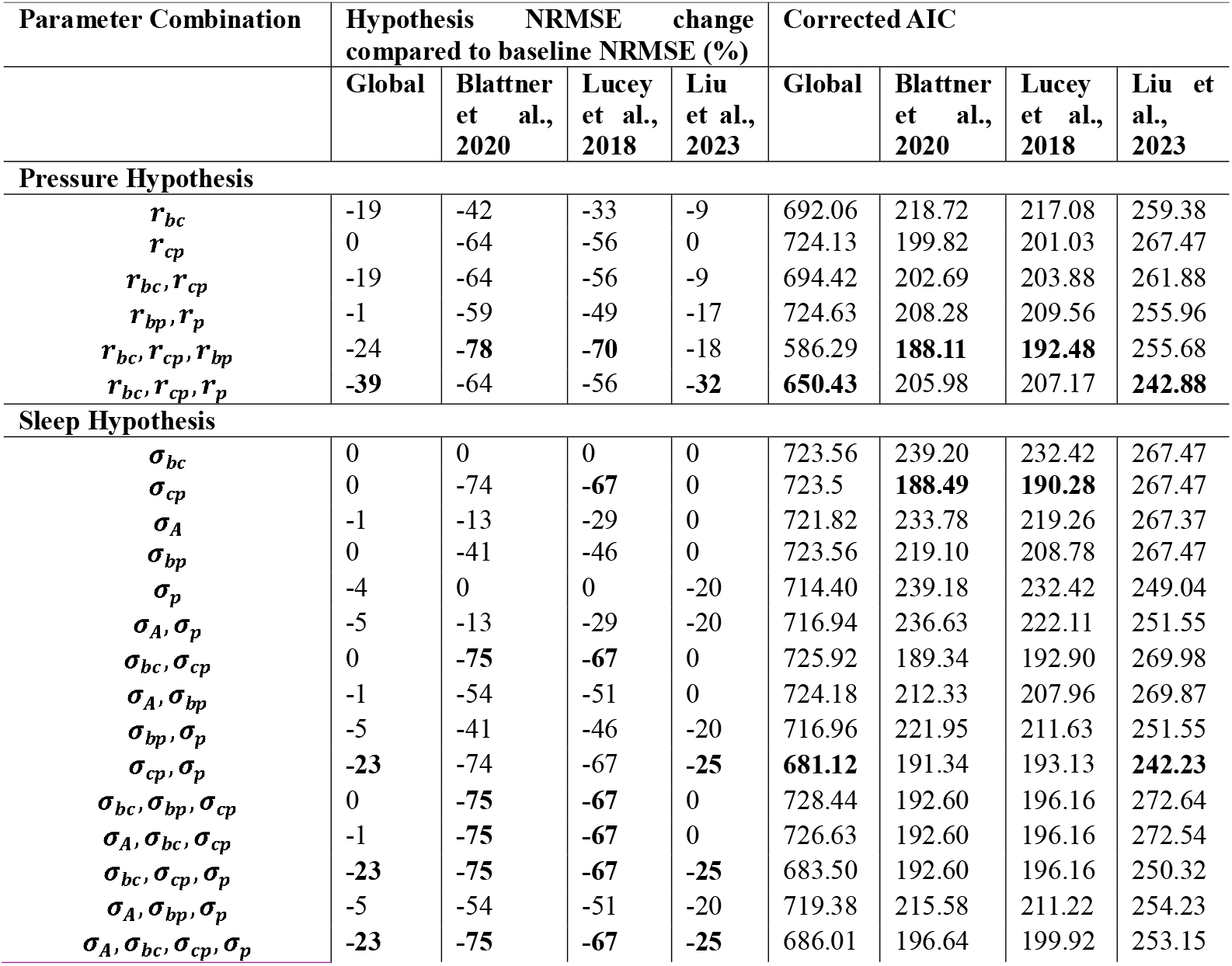
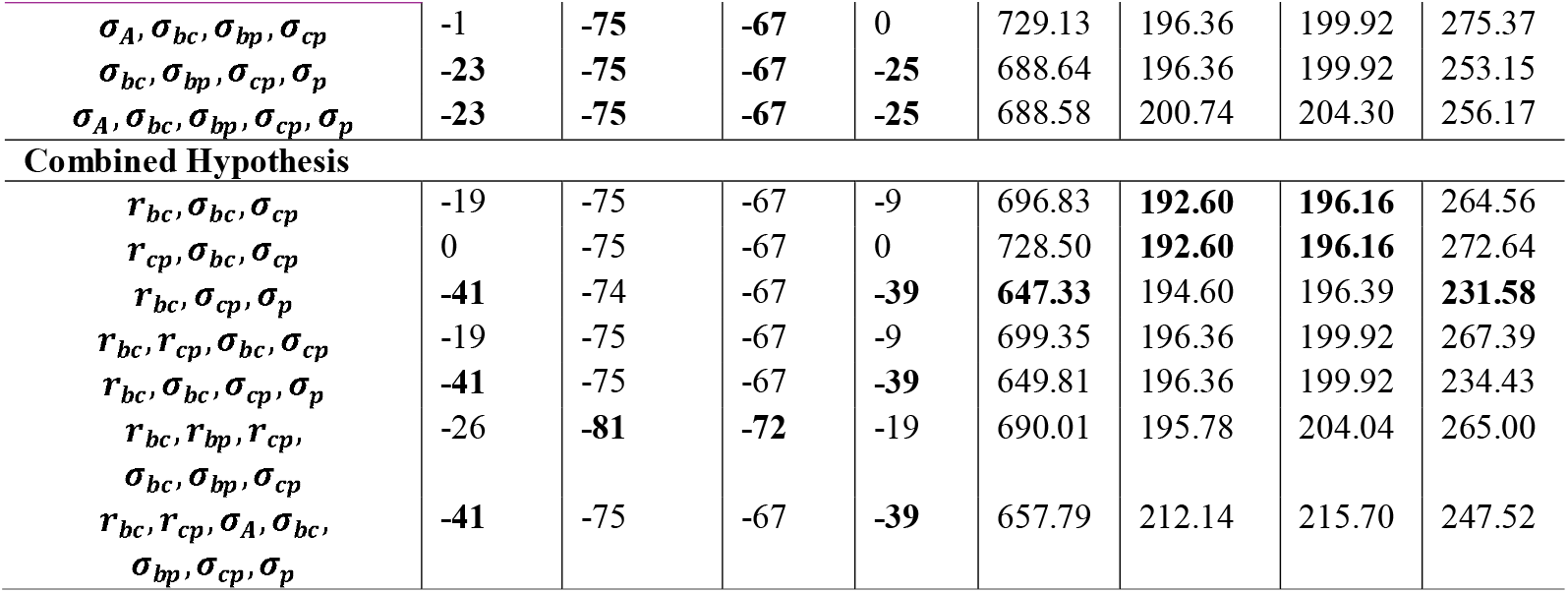
Changes in NRMSE and AICc for Lumbar Hypothesis Testing for Aβ42. Hypothesis NRMSE (percentage of baseline, %) represents the percentage change in fit quality compared to baseline NRMSE. For the global column, the baseline is the global fit across all datasets using default parameters. For individual dataset columns (Blattner et al., 2020, Lucey et al., 2018; Liu et al., 2023), the baseline is the respective individual fits from normal sleep-wake fitting. Negative values indicate the hypothesis improved the fit quality compared to baseline, positive values indicate worsened fit, and zero indicates no change. The lowest values for each dataset are highlighted in bold.

The hypotheses to find the effect of lumbar punctures on the Aβ dynamics were tested by fitting the model to the experimental data from the 36-hour sleep-wake protocols in Blattner et al.48, Lucey et al.4, and Liu et al.49, without detrending. This allowed us to identify the parameter set and values that best explain the observed increase in CSF Aβ42 concentration. In total, we evaluated 31 distinct parameter combinations.

#### Pressure Hypothesis

When assessing the performance of the model using corrected Akaike Information Criteria (AICc), all six tested parameter combinations, except for *r*_*cp*_, result in an improvement of the goodness of fit compared to the default set used for normal sleep-wake cycles, Table 2. The three-parameter model *r*_*bc*_,*r*_*cp*_,*r*_*p*_ achieved the lowest global AICc (model fit to all dataset simultaneously), indicating the best balance between fitting the data and model complexity. This fit was achieved by increasing *r*_*bc*_ and decreasing *r*_*cp*_,*r*_*p*_ compared to the default values, Table 3. Dynamically, these changes produced a steeper rise in CSF Aβ42 levels during wakefulness and slower decline during sleep, Figure 4. For individual studies, the best-fitting models varied. For Blattner et al.48 and Lucey et al.,4 the three-parameter model *r*_*bc*_, *r*_*cp*_,*r*_*bp*_ performed best, and for Liu et al.49, the three-parameter model *r*_*bc*_,*r*_*cp*_,*r*_*p*_ performed best.

**Table 3.** Best Model Parameter Values, NRMSE, and Corrected AIC (AICc) Corresponding to Different Hypotheses. The global fit represents parameter values optimised across all three datasets simultaneously (Blattner et al., 2020, Lucey et al., 2018, Liu et al., 2023). Individual dataset columns show parameters optimised for each study independently. Percentages in brackets indicate the relative change compared to the default parameter values for the normal sleep-wake cycles, with the sign demonstrating the direction of the change.

|  | Global Fit | Blattner et al., 2020 | Lucey et al., 2018 | Liu et al., 2023 |
| --- | --- | --- | --- | --- |
| **Pressure Hypothesis ($r_{bc}, r_{cp}, r_p$)** |  |  |  |  |
| $r_{bc}$ | 0.059 (55%) | 0.019 (0%) | 0.062 (0%) | 0.023 (53%) |
| $r_{cp}$ | $0.425 \times 10^{-2}$ (-21%) | 0.0102 (-34%) | 0.0113 (-28%) | $0.320 \times 10^{-2}$ (-0%) |
| $r_p$ | 0.361 (-15%) | 0.298 (0%) | 0.327 (9%) | 0.452 (-5%) |
| NRMSE | -39 | -64 | -56 | -32 |
| AICc | 650.43 | 205.98 | 207.17 | 242.88 |
| **Sleep Hypothesis ($\sigma_{cp}, \sigma_p$)** |  |  |  |  |
| $\sigma_{cp}$ | 2.861 (-53%) | 2.972 (-48%) | 2.242 (-45%) | 3.367 (-49%) |
| $\sigma_p$ | 2.519 (-41%) | 2.828 (-22%) | 2.272 (-43%) | 1.981 (-35%) |
| NRMSE | -23 | -74 | -67 | -25 |
| AICc | 681.12 | 191.34 | 193.13 | 242.23 |
| **Combined Hypothesis ($r_{bc}, \sigma_{cp}, \sigma_p$)** |  |  |  |  |
| $r_{bc}$ | 0.052 (37%) | 0.019 (0%) | 0.062 (0%) | 0.018 (20%) |
| $\sigma_{cp}$ | 3.733 (-39%) | 2.972 (-48%) | 2.242 (-45%) | 3.357 (-49%) |
| $\sigma_p$ | 2.922 (-31%) | 1.812 (-50%) | 2.137 (-47%) | 1.950 (-36%) |
| NRMSE | -41 | -74 | -67 | -39 |
| AICc | 647.33 | 194.60 | 196.39 | 231.58 |

#### Sleep Hypothesis

The two-parameter model *σ*_*cp*_, *σ*_*p*_ achieved the lowest global AICc, Table 2. This fit was achieved by decreasing both *σ*_*cp*_ and *σ*_*p*_ compared to baseline (Table 3). Dynamically, reduced *σ* parameters decreased overnight clearance, reproducing the experimental accumulation pattern for CSF Aβ42 (Figure 4). For the individual studies, different models perform best for different datasets, Table 2. The best model for Blattner et al.48 and Lucey et al.4, is the single parameter model *σ*_*cp*_ and for Liu et al.49, the best model was the two-parameter model *σ*_*cp*_, *σ*_*p*_.

#### Combined Pressure and Sleep Hypothesis

Based on the global AICc, the three-parameter model *r*_*bc*_, *σ*_*cp*_, *σ*_*p*_ achieved the lowest AICc, Table 2. This fit was achieved by the combined adjustment to the parameters with increasing *r*_*bc*_ and decreasing *σ*_*cp*_, *σ*_*p*_. Dynamically, the combined hypothesis reduced overnight clearance while enhancing the wake time influx into CSF, reproducing experimental dynamics (Figure 4a-d). For individual studies, three-parameter models performed best for Blattner et al.48, (*r*_*bc*_, *σ*_*bc*_, *σ*_*cp*_ and *r*_*cp*_,*σ*_*bc*_, *σ*_*cp*_ ) and Lucey et al.4, (*r*_*bc*_,*σ*_*bc*_, *σ*_*cp*_, and *r*_*cp*_,*σ*_*bc*_, *σ*_*cp*_), while for Liu et al.49, the three-parameter model (*r*_*bc*_, *σ*_*cp*_, *σ*_*p*_) performed best, Table 2.

Comparing the global AICc values, the combined hypothesis achieved the lowest AICc of 647.33 with ΔAICc of 3.10 relative to pressure hypothesis (AICc=650.43) and ΔAICc of 33.79 relative to the sleep hypothesis (AICc=681.12) (Table 2). Following Burnham & Anderson criteria72, ΔAICc of 3.10 indicates moderate rather than strong support, meaning the pressure and combined hypotheses cannot be definitely separated on statistical grounds alone. The sleep-only hypothesis receives considerably less support (ΔAICc>10). Given both pressure changes and some degree of sleep disruption are expected to co-occur during lumbar catheter protocols, the combined hypothesis remains the most plausible explanation for the observed CSF Aβ42 accumulation during lumbar puncture protocols. However, examining the individual study fits shows an important distinction in model consistency. The pressure hypothesis shows that the three-parameter model, *r*_*bc*_,*r*_*cp*_,*r*_*bp*_ performs best for Blattner et al.48, and Lucey et al.4, and the global optimal model (*r*_*bc*_,*r*_*cp*_,*r*_*p*_) is best for Liu et al.49. The sleep hypothesis shows that while single-parameter model *σ*_*cp*_ performs better for Blattner et al.48, and Lucey et al.4, the two-parameter model *σ*_*cp*_, *σ*_*p*_ performs better for Liu et al.49. The combined hypothesis shows a partial consistency with simpler three-parameter models (*r*_*bc*_, *σ*_*bc*_, *σ*_*cp*_ or *r*_*cp*_, *σ*_*bc*_, *σ*_*cp*_ ) performing better for Blattner et al.48 and Lucey et al.4, while the global optimal three-parameter model performs best for Liu et al.49. Given these mixed consistency patterns, which may reflect inter-individual differences in cerebrovascular architecture, CSF dynamics, and blood-CSF barrier functions73, we cannot definitively conclude that any single hypothesis model provides a superior explanation. However, the combined hypothesis offers a mechanistic understanding, and it is the most physiologically representative explanation, as both pressure-mediated transport changes and sleep-dependent clearance impairment are expected to operate simultaneously during the lumbar puncture protocols.

### Model Generalisability to Aβ40

The model successfully reproduces experimental Aβ40 dynamics across all three datasets, as shown in Figure 5a and 5b, with both isoforms exhibiting similar temporal dynamics. Using the same default parameter set with adjusted production (*A*_*Aβ*40_ = 8.87 \**A*_*Aβ*42_ ) and systemic clearance (*r*_*p,Aβ*40_ *= r*_*p,Aβ*42_ /1.44), the model maintains characteristic oscillations with brain Aβ40 concentrations declining from 2473 pgml^-l^ during wake to 2119 pgml^-l^ during sleep (∼14% decline). The CSF Aβ40 concentrations decline from 6950 pgml^-l^ to 5881 pgml^-l^ during sleep (∼13% decline) and the plasma compartment going from 233 pgml^-l^ to 197 pgml^-l^ during sleep (∼15% decline) (Figure S6).

**Figure 5.**
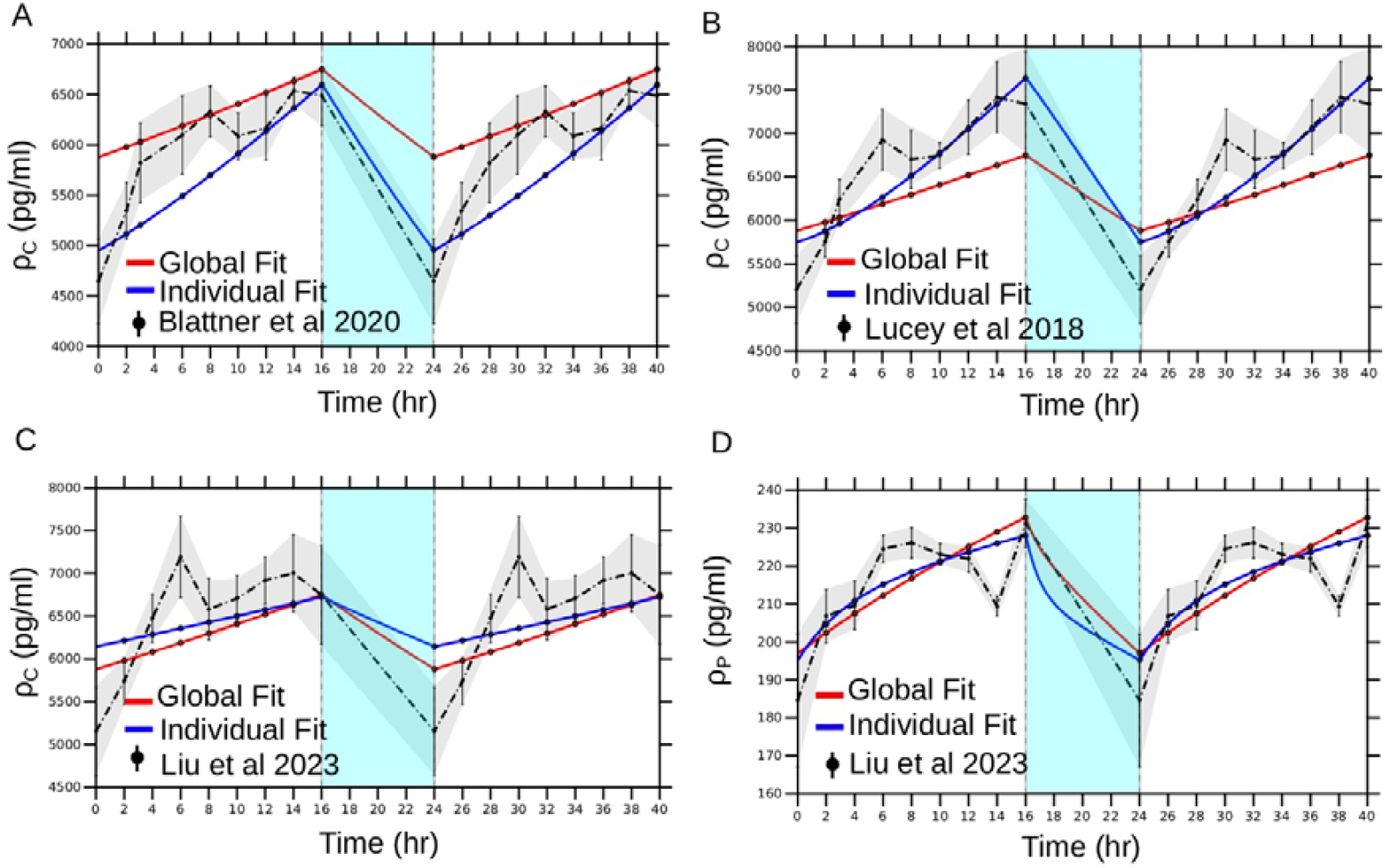
Model Fitting for Normal Sleep-Wake Cycle on Aβ40. **(A)** CSF data (Blattner et al., 2020)^13^ **(B)** CSF data (Lucey et al., 2018)^14^ **(C)** CSF data (Liu et al., 2023)^15^ **(D)** Plasma data (Liu et al., 2023)^15^. The grey shaded area denotes the experimental standard deviation. The cyan shaded area corresponds to the sleep state. The vertical dashed line at t=16 and t=24 denotes the onset and ending of the sleep state.

To test the mechanisms associated with the lumbar puncture effect on the dynamics of Aβ40, all three hypotheses (pressure hypothesis, sleep hypothesis, and combined hypothesis) were evaluated without any additional parameter optimisation. All hypothesis-specific parameters and their combinations remained identical to those fitted for Aβ42, and results are shown in Table S1 for completeness. The models that performed best in the Aβ42 scenario were also the best models for Aβ40 (Table 4), with the dynamics shown in Figure 6a-d.

**Table 4.** Model NRMSE and Normalised Corrected AIC (AICc) Corresponding to the Best Selected Hypotheses from Aβ42 Model Fitting and Applied to Aβ40 Data.

|  | Global Fit | Blattner et al., 2020 | Lucey et al., 2018 | Liu et al., 2023 |
| --- | --- | --- | --- | --- |
| **Pressure Hypothesis ($r_{bc}, r_{cp}, r_p$)** |  |  |  |  |
| $r_{bc}$ | 0.059 (55%) | 0.019 (0%) | 0.062 (0%) | 0.023 (53%) |
| $r_{cp}$ | $0.425 \times 10^{-2}$ (-21%) | 0.0102 (-34%) | 0.0113 (-28%) | $0.320 \times 10^{-2}$ (-0%) |
| $r_p$ | 0.361 (-15%) | 0.298 (0%) | 0.327 (9%) | 0.452 (-5%) |
| NRMSE | -37 | -62 | -43 | -30 |
| AICc | 1423.55 | 393.34 | 396.47 | 651.63 |
| **Sleep Hypothesis ($\sigma_{cp}, \sigma_p$)** |  |  |  |  |
| $\sigma_{cp}$ | 2.861 (-53%) | 2.972 (-48%) | 2.242 (-45%) | 3.367 (-49%) |
| $\sigma_p$ | 2.519 (-41%) | 2.828 (-22%) | 2.272 (-43%) | 1.981 (-35%) |
| **NRMSE** | -32 | -65 | -53 | -26 |
| **AICc** | 1421.46 | 390.15 | 393.27 | 649.06 |
| **Combined Hypothesis ($r_{bc}, \sigma_{cp}, \sigma_p$)** |  |  |  |  |
| $r_{bc}$ | 0.052 (37%) | 0.019 (0%) | 0.062 (0%) | 0.018 (20%) |
| $\sigma_{cp}$ | 3.733 (-39%) | 2.972 (-48%) | 2.242 (-45%) | 3.357 (-49%) |
| $\sigma_p$ | 2.922 (-31%) | 1.812 (-50%) | 2.137 (-47%) | 1.950 (-36%) |
| **NRMSE** | -39 | -65 | -53 | -33 |
| **AICc** | 1423.56 | 393.41 | 396.53 | 651.64 |

**Figure 6.**
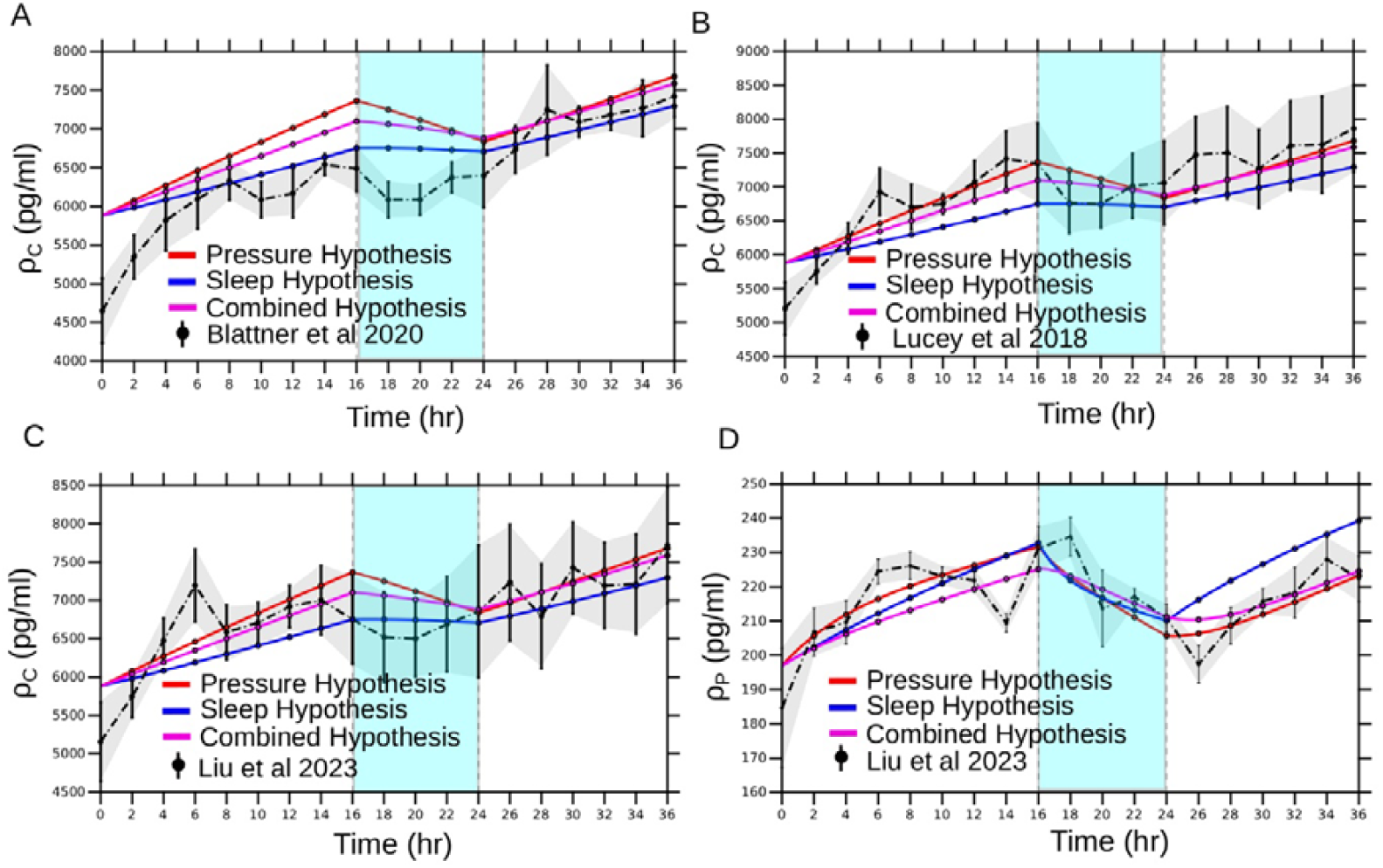
Model Fit to experimental Aβ40 Data. **(A)** CSF data (Blattner et al., 2020), **(B)** CSF data (Lucey et al., 2018), **(C)** CSF data (Liu et al., 2023), **(D)** Plasma data (Liu et al., 2023). The red curve corresponds to the pressure gradient hypothesis fit, the blue curve corresponds to the sleep disruption hypothesis fit, and the magenta curve corresponds to the combined hypothesis fit. The grey shaded area denotes the experimental standard deviation. The cyan shaded area corresponds to the sleep state. The vertical dashed line at t=16 and t=24 denotes the onset and ending of the sleep state.

## Discussion

Our three-compartment model simulates the dynamic interplay of Aβ production, transport, and clearance across the brain, CSF, and plasma in healthy individuals, with the sleep-wake cycle as a regulator. The key findings from the model are: (i) the model quantitatively reproduces experimental observations for CSF and plasma Aβ oscillations across sleep-wake cycles, (ii) the model predicts brain Aβ dynamics that remain experimentally inaccessible in humans, (iii) the model generalises to Aβ40 dynamics and (iv) the model provides mechanistic explanation for CSF Aβ accumulation during lumbar puncture protocols.

The model predicts Aβ42 dynamics across three compartments modulated by production and clearance rates. The brain compartment exhibits ∼15% change in concentration between wake to sleep transition consistent with long brain time constant (*τ*_*b*_ ≈ 14.8 hr during sleep) relative to 8 hr sleep duration. Although modest within a single night, the Aβ clearance represents a recurring mechanism where its disruption due to poor sleep quality or reduced sleep duration could lead to progressive Aβ accumulation over the years. CSF shows smaller relative oscillations (∼14%), compared to brain (∼15% variation) and plasma (∼24% variation), despite receiving a direct input from the brain, because sleep simultaneously increases brain-to-CSF influx (*σ*_*bc*_ *r*_*bc*_ ) and CSF-to-plasma efflux (*σ*_*cp*_ *r*_*cp*_). These competing fluxes balance each other, positioning CSF as a regulatory interface74.

Under lumbar puncture conditions using the combined hypothesis (*r*_*bc*_, *σ*_*cp*_, *σ*_*p*_), Aβ concentration in CSF during sleep is predicted to decline by only 3.1% (compared to 14% under normal sleep-wake cycles) which is consistent with the experimental datasets (Lucey et al.4, (∼2.4%) and Liu et al.49, (∼0.5%)). Blattner et al.48, shows an increase of 8.7% suggesting that the effect of indwelling lumbar catheter was more pronounced in the dataset. The plasma compartment shows a smaller absolute fluctuation (15-19 pgml^-l^, 4 pgml^-l^ range) compared to the brain (40 pgml^-l^ range) and CSF (98 pgml^-l^ range). This attenuation results from large plasma volume (∼5 L in plasma vs 0.15 L in CSF) and the filtering effect of the BBB transport20,74, which modulates the transmission of CNS Aβ42 fluctuation to systemic circulation. The combined hypothesis model (*r*_*bc*_,*σ*_*cp*_,*σ*_*p*_) predicts a ∼7.1% decline in Aβ42 plasma concentration from wake to sleep which is comparable to the ∼5.9% drop observed in Liu et al.49,. The agreement between the model predictions and experimental studies supports the model’s validity in capturing the sleep-associated Aβ clearance dynamics.

The model generalises to Aβ40 by adjusting only production rate and systemic clearance while keeping all other pathways identical to Aβ42. This suggests that sleep-wake regulation of Aβ dynamics operates through shared physiological mechanisms that do not discriminate between the Aβ isoforms. It should be noted that this finding is specific to amyloid-negative individuals, as deposition of Aβ into plaques in amyloid-positive participants would introduce isoform-specific dynamics that the present model does not capture.

While isoform-specific differences exist in LRP1 affinity, aggregation probability, and enzymatic degradation rates75, the dominant clearance mechanisms in amyloid-negative individuals are bulk flow processes including glymphatic clearance and CSF circulation76-78Because glymphatic and CSF based bulk flow mechanisms are thought to be primarily size-dependent, they are expected to be largely non-selective between closely related Aβ isoforms differing only by two amino acids79.

The stable concentration ratio of Aβ40:Aβ42 in CSF (∼8.87, Figure S7) reflects different production rates (Aβ40 produced ∼9 times more than Aβ42). The elevated ratio of Aβ40:Aβ42 in plasma (∼12.78, Figure S7) indicates slower peripheral clearance of Aβ40, consistent with experimental observation of faster Aβ42 fractional turnover61. Both isoforms display a comparable relative decline during sleep of ∼15-20% (model prediction), supporting the idea that central clearance pathways of brain-to-CSF and CSF-to-plasma, process both isoforms similarly through shared bulk flow mechanisms24.

The model’s ability to reproduce experimental Aβ42 and Aβ40 dynamics indicates that it captures the essential features of Aβ regulation along the sleep-wake cycles. The identified default parameter values are consistent with experimental observations and constraints:

– The baseline Aβ42 production rate (*A*=16.2 pgml^-l^h^-l^) is reduced by ∼23% during sleep (*σ*_*A*_ = 0.772), consistent with experimental observations reporting a 10-30% decrease in neuronal activity and Aβ levels during sleep50. This decrease in production, combined with sleep-dependent clearance, produces a ∼15% oscillation in brain Aβ42 concentration across the sleep-wake cycle. Aβ levels increase during wakefulness with higher neuronal firing and metabolism80 and decrease during SWS when neuronal activity declines81.
– The total CNS clearance rate (*r*_*bc*_ *+ r*_*bp*_ *+ r*_*cp*_ ≈ 0.057 h^-l^) is comparable to the in vivo measurements (0.083 h^-l^) indicating the model correctly balances the brain production against total clearance. The BBB contribution of ∼26-37% of brain efflux (Figure S10) aligns with Roberts et al.74, reporting a ∼25% clearance of Aβ via BBB pathways, suggesting that the majority of brain Aβ clearance occurs via brain-to-CSF (*r*_*bc*_,*σ*_*bc*_ ) and CSF-to-plasma (*r*_*cp*_, *σ*_*cp*_ ) routes.
– Systemic plasma clearance rate was obtained as *r*_*p*_ = 0.427 h^-l^, which corresponds to a plasma Aβ42 half-life of ∼1.62 h. This is faster than the experimentally observed 2-3 h half-life61 but remains within the feasible physiological range. This difference likely arises because *r*_*p*_ is a net systemic clearance parameter which absorbs contributions from renal filtration, hepatic metabolism and peripheral tissue uptake simultaneously. Overall, the globally fitted parameters offer a physiologically consistent representation of both production and clearance pathways, balancing available experimental data with established theoretical constraints.

Some of the fitted model parameters show differences from published experimental values due to methodological differences. For example, the model’s BBB clearance rate (*r*_*bp =*_ 0.014 h^-l^) is lower than Roberts et al’s.74, 0.140 h^-l^ likely because experimental data measures total soluble Aβ clearance (Aβ38, Aβ40, Aβ42 combined) while our model is isoform specific. Similarly, the model’s CSF-to-plasma clearance (*σ*_*cp*_ *r*_*cp*_ ) differs from Bateman et al’s51 by approximately two-fold, reflecting the same isoform-specific versus total Aβ comparison. Despite these individual parameter differences, the model successfully captures 24-hour Aβ42 oscillations across multiple datasets, indicating that the underlying physiological relationships are appropriately represented.

Prior modelling approaches to Aβ dynamics have addressed different aspects of the Aβ production, clearance, and aggregation, often operating on different spatial and temporal scales40,42-44,46. These models have deepened our understanding of pathological mechanisms, but most of them do not incorporate sleep-wake cycle regulation. Our model addresses this by explicitly representing how sleep-wake transitions alter production and clearance rates, enabling the simulation of 24-hour Aβ oscillations observed experimentally in CSF and plasma. These predictions extend to brain dynamics, which are important because they represent the compartments where pathological accumulation originates, yet experimental access requires invasive microdialysis, which is not feasible in human studies82.

Recent model by Dagum et al.47 is most similar to our model in that it also considers sleep and wake states. Dagum et al. model has six compartments and predicts how sleep-associated changes in Aβ and tau release and glymphatic clearance affect morning plasma biomarker levels. In a randomised crossover trial, they demonstrated that neurophysiological markers of glymphatic function strongly predicted morning plasma Aβ and tau levels and that clearance is enhanced during normal sleep compared to sleep deprivation. While both models (Dagum et al. and ours) incorporate sleep-wake regulation, they differ in structure and objective. Dagum et al’s47 model includes local degradation and aggregate compartments with literature-derived parameters. These processes are negligible for cognitively normal, amyloid-negative individuals over 24–36-hour timescales and are therefore absorbed into net clearance parameters in our model. Also, Dagum et al47 modulate only glymphatic exchange and cellular release during sleep, whereas our model extends sleep-state modulation to all clearance pathways simultaneously, and is fitted globally against the published CSF and plasma timeseries data. The two models are therefore complementary: Dagum et al. addresses the mechanistic basis of biomarker release during sleep and sleep deprivation; our model accounts for 24-hour Aβ oscillations under heathy physiological conditions against which sleep deprivation and long-term Aβ accumulation can be simulated and quantified.

Experimental data using lumbar puncture for CSF collection show a day-to-day increase in CSF amyloid concentration (Figure S5). A consistent pattern is found across all three hypotheses. The brain-to-CSF (*r*_*bc*_ ), CSF-to-plasma (*r*_*cp*_, *σ*_*cp*_ ) and systemic clearance (*r*_*p*_, *σ*_*p*_) pathways appear as common parameters across the best model in each hypothesis. The best model in the pressure hypothesis involves *r*_*bc*_,*r*_*cp*_ and *r*_*p*_, the best model of sleep hypothesis involves *σ*_*cp*_ and *σ*_*p*_ and the best model of the combined hypothesis involves *r*_*bc*_,*σ*_*cp*_ and *σ*_*p*_. The repeated appearance of brain-to-CSF, CSF-to-plasma and systemic clearance suggests that the model predicts these pathways as the primary sites where lumbar puncture protocols may disrupt Aβ homeostasis, whether through altered pressure gradient affecting clearance dynamics or sleep disruption affecting the glymphatic or blood-CSF barrier function. The brain-to-CSF pathway (*r*_*bc*_ ) appears only in the model’s pressure and combined hypotheses, which is consistent with pressure-mediated effects at the brain-CSF interface, while sleep disruption is inferred to primarily affect CSF-to-plasma and systemic clearance. This represents an important prediction from the model suggesting that the lumbar puncture-associated pressure changes and sleep disruption do not perturb Aβ homeostasis through the same clearance steps. Instead, they may act on different segments of the clearance pathways. Pressure changes from the CSF collection during lumbar puncture may alter the upstream step driving more Aβ from brain ISF to CSF. Sleep disruption may thus impair the downstream clearance of Aβ from CSF into plasma and systemic circulation. The combined hypothesis offers a more mechanistic advantage by acknowledging that both processes may operate simultaneously. It shows the lowest global AICc among the tested hypotheses and the biological plausibility of both mechanisms is well supported.

Studies have shown that high-frequency lumbar CSF sampling, such as every 30 minutes, significantly elevates Aβ levels compared to low-frequency protocols (e.g., every 4 hours), which maintains more stable concentrations, suggesting that sampling intensity disrupts clearance or redistributes CSF Aβ concentration gradients83. The magnitude of this linear rise has been shown to depend on the frequency and volume of amount of CSF withdrawn, most likely through redistribution of CSF flow towards the lumbar space84, consistent with the pressure hypothesis explored with the model. This supports the hypothesis that frequent sampling perturbs physiological Aβ dynamics, possibly by altering rostral-caudal gradients or clearance efficiency85. It is important to note that this sampling induced rise in Aβ levels is specific to amyloid-negative individuals and is attenuated in amyloid-positive individuals, while Aβ40 is unaffected suggesting that amyloid plaques act as a concentration sink and isolates soluble Aβ42 from the CSF pool and dampens the catheter-induced rise84. Moreover, multiple recent studies have demonstrated a direct connection between poor sleep quality and reduced glymphatic efficiency, providing support for the view that restorative sleep is crucial for optimal brain clearance86,87. Poor subjective sleep quality and shorter sleep duration correlate with decreased glymphatic function and cognitive decline, reinforcing the importance of sleep in maintaining efficient clearance of Aβ and tau proteins87,88. All three hypotheses successfully generalised to Aβ40 dynamics without additional parameter optimisation.

Our model makes a prediction for how lumbar puncture protocols create measurement artifacts. Model fitting identifies brain-to-CSF (*r*_*bc*_ ), CSF-to-plasma (*r*_*cp*_,*σ*_*cp*_ ) and systemic clearance (*r*_*p*_,*σ*_*p*_) pathways as requiring adjustments to reproduce lumbar puncture effects when compared to the normal sleep-wake cycles, suggesting these pathways might be affected by experimental protocols. The 2-hour sampling intervals used in the studies (Blattner et al.48, Lucey et al.4, Liu et al.49), show sustained Aβ elevation in CSF on day 2 compared to day 1 (Figure S5). This suggests that non-invasive approaches, such as MRI-based glymphatic imaging89, may provide more stable measurements for temporal studies by avoiding the procedural effects introduced by repeated CSF withdrawals. However, MRI-based methods have their own limitations, such as measuring surrogate markers of clearance function, such as contrast tracer movements, rather than direct Aβ dynamics90, dependence on exogenous tracer administration with its own pharmacokinetic complexities91,92, and limited temporal resolution93,94. These limitations mean that MRI-based glymphatic imaging complements rather than replaces lumbar catheter approaches and future studies would ideally combine both approaches to cross validate clearance estimates. Complementary approaches such as sleep EEG monitoring offer an additional approach where continuous EEG recording during sleep-wake cycles capture slow-wave activity95. Also, non-invasive devices, such as used by Dagum et al.86, offers a promising method to capture sleep-active glymphatic functions. For studies specifically investigating the effect of sleep on Aβ clearance, the model’s sensitivity to sleep-dependent clearance parameters (Figure S1-S3) indicates that any procedural disruption to sleep quality, including body position96, could affect experimental interpretation. A formal global sensitivity analysis confirmed that baseline clearance rates and production amplitude are the dominant drivers of variance across all three compartments, with sleep-dependent scaling parameters showing comparatively lower sensitivity (Fig. S3). This suggests that the experimental designs must control for this artifact to isolate the biological effect of interest.

Overall, our compartmental model has successfully predicted amyloid-beta (Aβ) sleep-wake dynamics across the brain and reproduced the Aβ dynamics, within experimental levels, in CSF and plasma compartments. The model reveals that lumbar puncture protocols create measurement artifacts with CSF-to-plasma and systemic clearance affected the most. This work provides a minimal model framework that can be used for investigating pathological states and exploring the role of sleep-wake cycles on the Aβ dynamics.

### Limitations and Future Extensions

Despite successfully reproducing experimental data, the model is a significant simplification of reality, which introduces limitations to its use. We represent sleep and wake as discrete binary states and multiple biological processes, such as enzymatic degradation, phagocytosis, and active transport, are represented as a single effective rate constant. Thus, fitted parameters represent overall effective values that reproduce the observed dynamics rather than direct measurements of isolated mechanisms. The model does not have a single global minimum when fitted to experimental data; instead, a number of (similar) parameter sets describe the dynamics (parameter identifiability analysis shown in Figs S1-S3 and Table S2). This should be addressed in the future when more experimental data become available.

A useful future application of the model would be to predict brain dynamics from plasma biomarkers, e.g., to track drug effects. However, the current model will need further validation on new data under various experimental protocols to enable such functionality. This is because significant differences in brain and CSF compartments may produce only minor (experimentally indistinguishable) changes in plasma and, vice versa, significant changes in plasma may be due to peripheral clearance rather than changes in brain or CSF (examples in Figs S8, S9).

The unidirectional clearance framework is appropriate for healthy homeostasis but excludes bidirectional BBB transport relevant in pathological states46. We do not model CSF volume dynamics when simulating lumbar puncture effects. Concentration changes are associated with altered transport kinetics and sleep disruption, but repeated CSF withdrawals also lead to CSF volume depletion, which might affect the concentration of Aβ. Future model refinement could address these limitations. The model does not account for the delay between brain CSF and CSF in the lumbar region. Future extensions could incorporate an explicit transport delay to characterise brain and lumbar CSF dynamics separately.

Modelling circadian regulation, beyond sleep-wake states, would more accurately represent the glymphatic fluctuations. This is particularly relevant for studying irregular sleep patterns, such as in shift workers, who show increased dementia risk potentially linked to altered clearance97, individuals with sleep disorders, or older adults in whom age-related deterioration results in changes in sleep physiology98. The model can be extended to incorporate bidirectional BBB transport, as done by Ficiara et al.46, to simulate diseased conditions such as Alzheimer’s Disease.

Similarly, the model could be extended to include additional compartments representing cellular uptake, local degradation, and oligomerisation dynamics, as implemented by Dagum et al.47, to enable mechanistic differentiation between production-driven versus clearance-driven pathological changes through biomarker ratio analysis. Further, extending the timescale from hours to months and years would facilitate the investigation of cumulative effects, such as how chronic sleep disruption translates acute clearance disruption into long-term Aβ accumulation. These extensions would broaden the model’s utility from healthy state baseline dynamics towards exploring disease progression.

## Resource Availability

### Lead Contact

Further information and requests should be directed to and will be fulfilled by Satyam Sangeet

### Materials Availability

This study did not generate any new material

### Data and Code Availability

All the data for model fitting were taken from published research articles. The code for model fitting, simulation, hypothesis testing are available at https://github.com/satyamsangeet/Amyloid_Clearance_3C

### Author Contributions

Conceptualization, S.P. and S.S; Methodology, S.S. and S.P.; Investigation, S.S. and S.P.; Visualisation, S.S.; Writing – original draft, S.S. and S.P..; Writing – review and editing, S.S., C.L.P., C.M.H., A.L.D., B.P.L. and S.P; Supervision, S.P.

## Acknowledgements

This work was supported by the Australian Research Council (ARC DP230101113), Faculty of Science Research Scholarship from University of Sydney, and a top up scholarship from the Centre for Chronic Diseases of Ageing (CCDA), Woolcock Institute of Medical Research.

## Methods

### Experimental Data and Model Fitting

#### Experimental Data for Temporal Aβ Dynamics

To ensure an accurate and physiologically relevant temporal model, we selected experimental studies that met the following criteria: 1) Participants were cognitively healthy adults, 2) Sampling was performed in a time-dependent manner across at least 36 hours, during normal sleep-wake cycle. Three studies satisfied this criterion: Blattner et al.48, Lucey et al.^4^, and Liu et al.49, with Blattner et al.48 and Lucey et al.4 exclusively reporting CSF Aβ levels and Liu et al.49 reporting simultaneous CSF and Plasma Aβ measurements. The selected studies reported serial Aβ42 measurements for CSF and/or plasma every 2 hours over the 36-hour experimental protocol. CSF and blood were collected via indwelling lumbar and intravenous catheters. Blattner et al.48 reported Aβ42 concentrations as percentage change from the mean value across the full 36-hour sampling period while Lucey et al.4 and Liu et al.49 reported Aβ42 concentrations as percentage change of baseline with Aβ42 normalised to the mean value from 07:00-19:00 (12 hours) prior to intervention (Table 5).

**Table 5.**
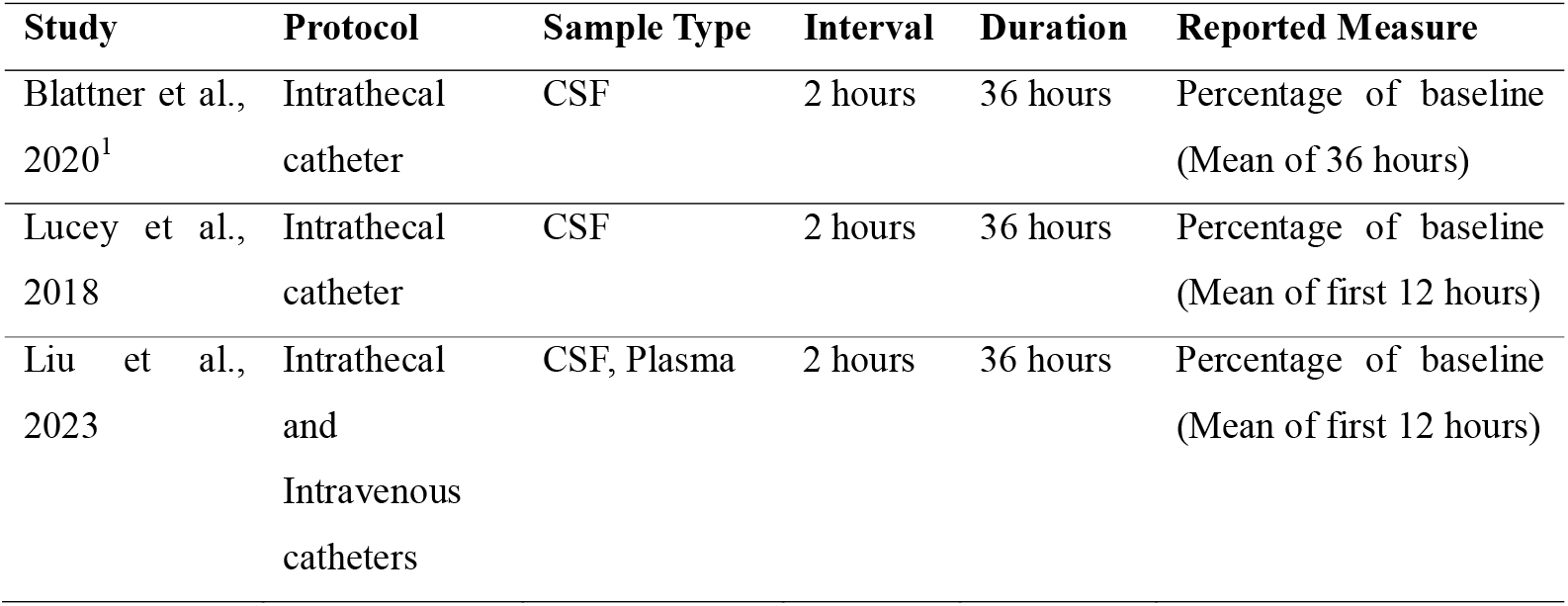
Experimental Protocol for the Collected Temporal Studies for Model Fitting.

| Study | Protocol | Sample Type | Interval | Duration | Reported Measure |
| --- | --- | --- | --- | --- | --- |
| Blattner et al., 2020<sup>1</sup> | Intrathecal catheter | CSF | 2 hours | 36 hours | Percentage of baseline (Mean of 36 hours) |
| Lucey et al., 2018 | Intrathecal catheter | CSF | 2 hours | 36 hours | Percentage of baseline (Mean of first 12 hours) |
| Liu et al., 2023 | Intrathecal and Intravenous catheters | CSF, Plasma | 2 hours | 36 hours | Percentage of baseline (Mean of first 12 hours) |

#### Experimental Data for Mean Physiological Aβ Levels in CSF and Plasma

To enable direct comparison and model fitting, we converted the reported percentage changes in Aβ42 concentration to absolute values (pg/ml) by multiplying each time point’s percentage by the reference mean baseline concentration. To determine the reference mean baseline concentration, we collected experimental data on Aβ concentrations in CSF and plasma from studies involving cognitively healthy participants (Table S3). Because of the variability in assay techniques and reporting formats across studies, we used strict inclusion criteria to ensure that the data were comparable and reliable. Specifically, we selected studies that employed the enzyme-linked immunosorbent assay (ELISA) method for Aβ quantification. Studies utilising other assay platforms were excluded or converted to ELISA-equivalent using validated conversion factors. The selected studies reported Aβ concentrations using diverse statistical measures, including means with standard deviations, median with interquartile ranges, and coefficient of variation. To provide a meaningful comparison, all the reported values were converted into a common format of mean ± standard deviation (SD). For studies reporting medians and percentile ranges, we applied statistical methods to estimate means and SDs under the assumption of normal distribution. When only ranges or coefficients of variation were available, we used standard formulas to approximate the SDs. To obtain average Aβ concentrations in CSF and plasma, we used the decomposition of the Mean and Standard Deviation method. This method allowed us to aggregate data from multiple studies by weighting means and variances according to sample sizes. The resulting mean and SDs serve as baseline physiological levels of Aβ in CSF and plasma compartments, which are essential for model calibration and for converting the experimental measurements into absolute concentration units. Since there is an inherent variability in physiological Aβ levels due to individual differences, we calculated the theoretical concentration ranges using the interquartile range (IQR) method. This allowed us to calculate the lower and upper bounds for Aβ concentrations in CSF and plasma, which are used as constraints during model parameter estimation. This allowed us to standardise all temporal profiles to the same units, ensuring that model outputs could be directly validated against relevant concentration ranges. The relevant details of the experimental data are provided in Table S3.

#### Reconstruction of Aβ dynamics for Normal Sleep-Wake Cycle

Aβ concentration in brain interstitial fluids, CSF, and plasma compartments has been shown to fluctuate in a diurnal pattern. Experimental studies in both humans and mice demonstrate that Aβ levels rise during wakefulness and decrease during sleep80,99. While the literature does not specifically state that Aβ returns to precisely the same baseline at the start of every day, these studies consistently show the dynamic regulation of Aβ levels99. Aβ decreases with sustained sleep and tends to be restored towards lower, homeostatic levels following periods of prolonged wakefulness or sleep deprivation100.

However, capturing these patterns experimentally is challenging82. CSF sampling by lumbar puncture introduces a methodological issue where serial CSF collection produces an upward drift in measured Aβ concentration over time67,68. Experimental evidence has shown that the linear rise of Aβ is a function of the frequency and volume of CSF collection84. This upwards drift pattern is sometimes removed from reported data by de-trending85. This upwards drift is not observed in plasma, likely due to much higher plasma volume (∼5 L) compared to CSF (∼150 mL). In the experimental dataset used for model fitting4,48,49, CSF Aβ concentrations post sleep phase remained elevated compared to baseline (Figure S5), consistent with findings shown by Xu et al.,101 where in monkeys, lumbar punctures caused a rapid, sustained increase in CSF Aβ and tau for up to 10 days, likely due to disrupted CSF outflow and fluid shifting from the frequent CSF collections. To minimise these experimental artifacts and approximate the physiological Aβ dynamics for model fitting, we reconstructed the Aβ time series for a normal sleep-wake cycle using the first 16 hours of experimental data. Specifically, the first 16-hour wake period concentration profile was obtained and repeated for day 2 with intervening 8 hours sleep period containing no experimental data, resulting in a 40-hour timeseries.

#### Model Fitting

Normalised Root Mean Square Error (NRMSE) was used as the model optimisation objective function. To achieve reliable parameter estimation, MATLAB’s interior-point technique (*fmincon*)102 was used for model calibration. Strict convergence requirements were applied (maximum iterations: 1000, function evaluations: 5,000). To prevent convergence to local minima, a multi-start technique was used (n=20), with a varying initial condition sampled from physiologically relevant ranges with a fixed random seed to ensure reproducibility. The algorithm explored the parameter space within the biophysical constraint bounds until the change in the objective function fell below the function tolerance or the maximum iteration limit was reached. For model calibration, two fitting strategies were employed: a global fit and individual fits. In the global fit, a single parameter set was optimised to minimise the average error across all datasets using the mean of the four individual NRMSE values as the objective function. The global error was calculated as:

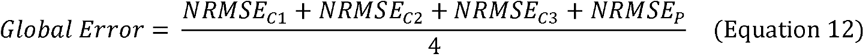

Where *NRMSE*_*Cl*_,*NRMSE*_*C2*_ and *NRMSE*_*C3*_ represents the NRMSE for CSF data from Blattner et al.48, Lucey et al.4, and Liu et al.49, respectively, and *NRMSE*_*P*_ is the NRMSE for plasma data from Liu et al.,49. For individual fits, model parameters were optimised separately for each dataset. In the case of Liu et al.49, which provided both CSF and plasma data, the individual error was calculated as the average of CSF and plasma NRMSE values:

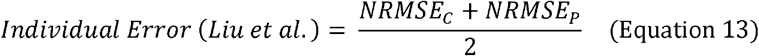

Where *NRMSE*_*C*_ and *NRMSE*_*P*_ are the NRMSE for CSF and plasma, respectively. For Blattner et al.48, and Lucey et al.4, the NRMSE was calculated for CSF data alone. The solution with minimal NRMSE was selected as the final fit.

We applied physiological constraints based on brain compartment clearance time constants to prevent unrealistic steady-state approaches during sleep-wake cycles. The nonlinear constraints (Equations 9 and 10) were implemented during optimisation, constraining the sum of transfer rates rather than individual pathways. To further prevent the brain compartment from reaching theoretical steady-state values due to unrealistic parameter combinations, we implemented a continuous penalty function. Since the brain’s regular sleep-wake cycles are essential to a healthy Aβ42 metabolism, the brain never functions at a steady state for Aβ42 concentrations in healthy physiology. The steady state concentration of brain Aβ42 during wake and sleep states is defined according to equation 4 and equation 5. Relative deviation of simulated brain concentration (*ρ*_*b*_*(t)*) from steady state during wake and sleep was quantified with a tolerance threshold (*δ* = 0.10) to define when concentrations are considered close to steady state. A closeness score was calculated for each time point, and the raw penalty (*P*_*r*_) was defined as the mean squared closeness across wake and sleep periods. To ensure proportional weighting (*P*_*w*_), the penalty was auto-calibrated relative to initial fitting error (NRMSE). The final objective function combined normalised data error and steady-state penalty:

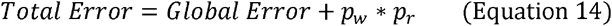

This error acts as a physiological regularisation term, restricting the parameter space to regions that produce realistic oscillatory Aβ42 brain dynamics.

### Effect of Lumbar Puncture on elevated CSF Aβ levels

To investigate the mechanistic contribution of experimental effect (lumbar puncture) on CSF Aβ42 levels, we formulated three primary hypotheses: 1) pressure-driven effects, 2) sleep-dependent clearance, and 3) combined mechanism incorporating both pathways. For each hypothesis, we identified and varied the specific model parameters associated with the relevant physiological process while maintaining all other parameters at their default values previously optimised against reconstructed data. Model fitting was performed on the full 36-hour time series of Aβ42 concentrations. A similar fitting protocol was followed as above, with NRMSE as the model optimisation function. The resulting best-fit parameters for each hypothesis were then used to assess the possibility of pressure, sleep, or a combined effect in accounting for the observed CSF Aβ levels.

### Model selection using Corrected Akaike Information Criteria (AICc)

To objectively compare the hypotheses explaining the CSF Aβ42 dynamics, we evaluated the model performance using the corrected Akaike Information Criteria (AICc), which balances the model fit and complexity. For model fit to the Liu dataset (consisting of both CSF and plasma experimental data) and the global fit (model fit to all datasets simultaneously), we employed the heteroscedastic likelihood framework where each compartment has its own variance parameter103. The log-likelihood for each compartment *j* is:

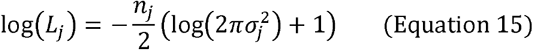

Where *n*_*j*_ is the number of observations in the compartment *j* and 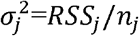 is the maximum likelihood estimate of variance for that compartment with *RSS*_*j*_ being the residual sum of squares. The total log-likelihood is the sum across all compartments

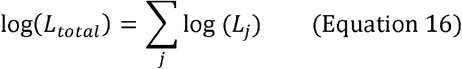

The AIC and corrected AIC (AICc) were calculated as:

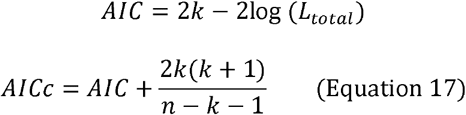

Where *k* is the total number of parameters (model parameters plus variance parameter) and *n* is the total number of observations across all compartments. This approach is used because the CSF and plasma compartments have different experimental measurement scales (∼700-800 pgml^-1^ for CSF vs 15-20 pgml^-1^ for plasma), resulting in different variances of the compartments, which would otherwise result in inflated AICc values if a homoscedastic approach (assuming the same variance for every compartment) is used. AICc values are interpretable only for comparing models fitted to the same dataset. Because of differences in sample sizes and variances of the datasets, AICc values obtained from different datasets are not directly comparable. The model with the lowest AICc value was selected as the best-fitting model.

### Model generalisability for Amyloid-β 40 (Aβ40)

To evaluate the generalisability of the model across Aβ species, we extend the baseline and hypothesis-specific model, originally fit to Aβ42, to predict Aβ40 dynamics. Importantly, no additional parameter fitting was performed. Aβ40 experimental concentrations were processed using the same pipeline as described above for Aβ42. Reported percentage changes of Aβ40 were converted to absolute concentrations (pgml^-l^) using reference baseline levels, which were derived from the published studies in cognitively healthy participants (see Supplementary). Reported measures were normalised to mean ± SD format to allow comparison across studies. The resulting baseline Aβ40 levels in CSF and plasma were used to calculate the concentration ratio (*ρ*_*Aβ*40_ : *ρ*_*Aβ*42_ ). Across the datasets, the average CSF ratio was ∼8.87, while the plasma ratio was higher at ∼12.78 (Figure S7). If the clearance process were identical for both species, the ratio would remain constant between CSF and plasma. The observed enrichment in plasma indicates a species-specific divergence in clearance or influx mechanisms. Two mechanistic scenarios were considered: 1) Slower clearance of Aβ40 from plasma to systemic circulation (*r*_*p*_) and 2) increased influx (*r*_*bp*_,*r*_*cp*_) of Aβ40 into the plasma relative to Aβ42. While both scenarios are biologically plausible, we adopted the first approach for methodological reasons. Implementing increased influx (scenario 2) would require adjusting brain-to-plasma (*r*_*bp*_) or CSF-to-plasma (*r*_*cp*_ ) transfer rates, which would alter upstream CSF Aβ40 concentrations according to steady state equations. This would disrupt the experimentally observed CSF ratio of 8.87 and compromise the model’s ability to simultaneously predict both compartments accurately. On the other hand, modulating plasma clearance (*r*_*p*_) provides a minimal adjustment that preserves CSF dynamics while accounting for the observed plasma enrichment. While this method allows us to test the model generalisability without complicating the predictions for upstream compartments, it does not rule out the possibility that differential influx processes could potentially be responsible for the observed concentration ratio (*ρ*_*Aβ*40_ : *ρ*_*Aβ*42_ ) differences.

To quantify this effect, we defined a plasma enrichment factor as the ratio of plasma-to-CSF enrichment:

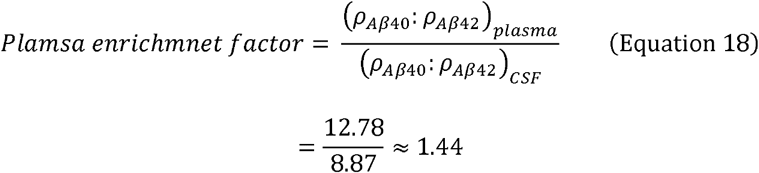

This factor was applied to plasma clearance rate by scaling *r*_*p*_ *→ r*_*p*_/1.44, thereby reducing the Aβ40 clearance relative to Aβ42. Production rate was simultaneously scaled by 8.87 (*A* → *A* * 8.87) to maintain the experimentally observed *ρ*_*Aβ*40_ : *ρ*_*Aβ*42_ ratio in CSF.

