## Supplementary File for "Quantitative Modelling of Amyloid-β Dynamics in Brain, CSF, and Plasma During Sleep and Wakefulness"

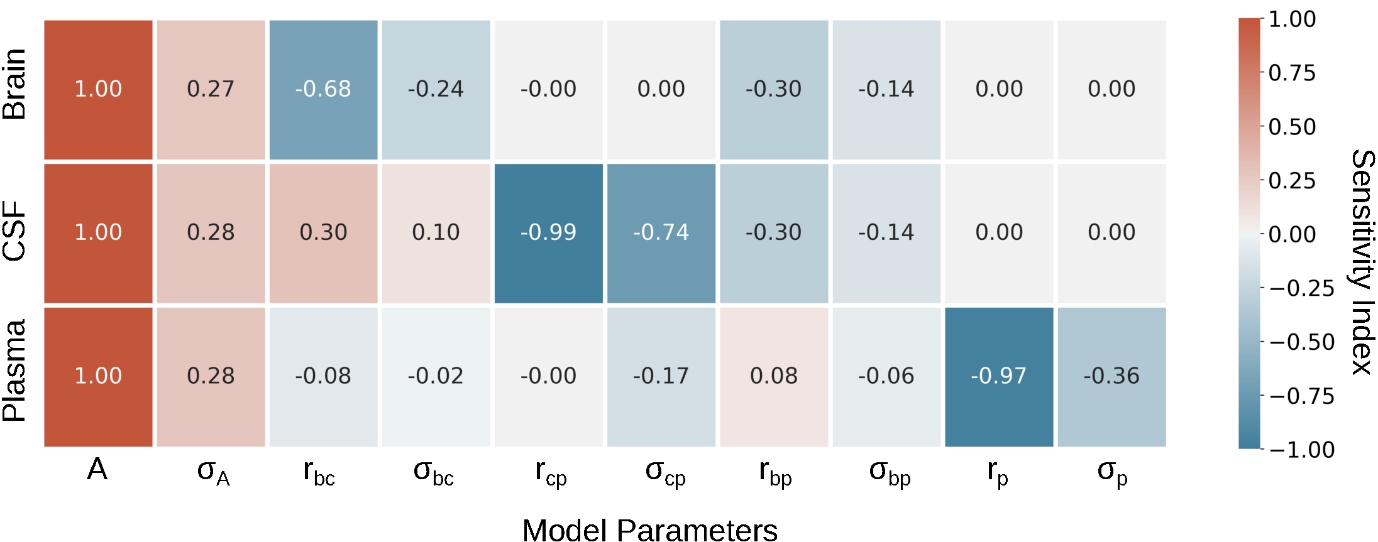

**Figure S1: Sensitivity Analysis of the model.** Local sensitivity indices for model parameters across three compartments. The rows correspond to the three compartments in the model, and the columns represent the analysed parameters. Sensitivity indices range from -1 to 1, as indicated by the colour bar, with positive values denoting a direct relationship and negative values indicating an inverse relationship between the parameter and the compartment output.

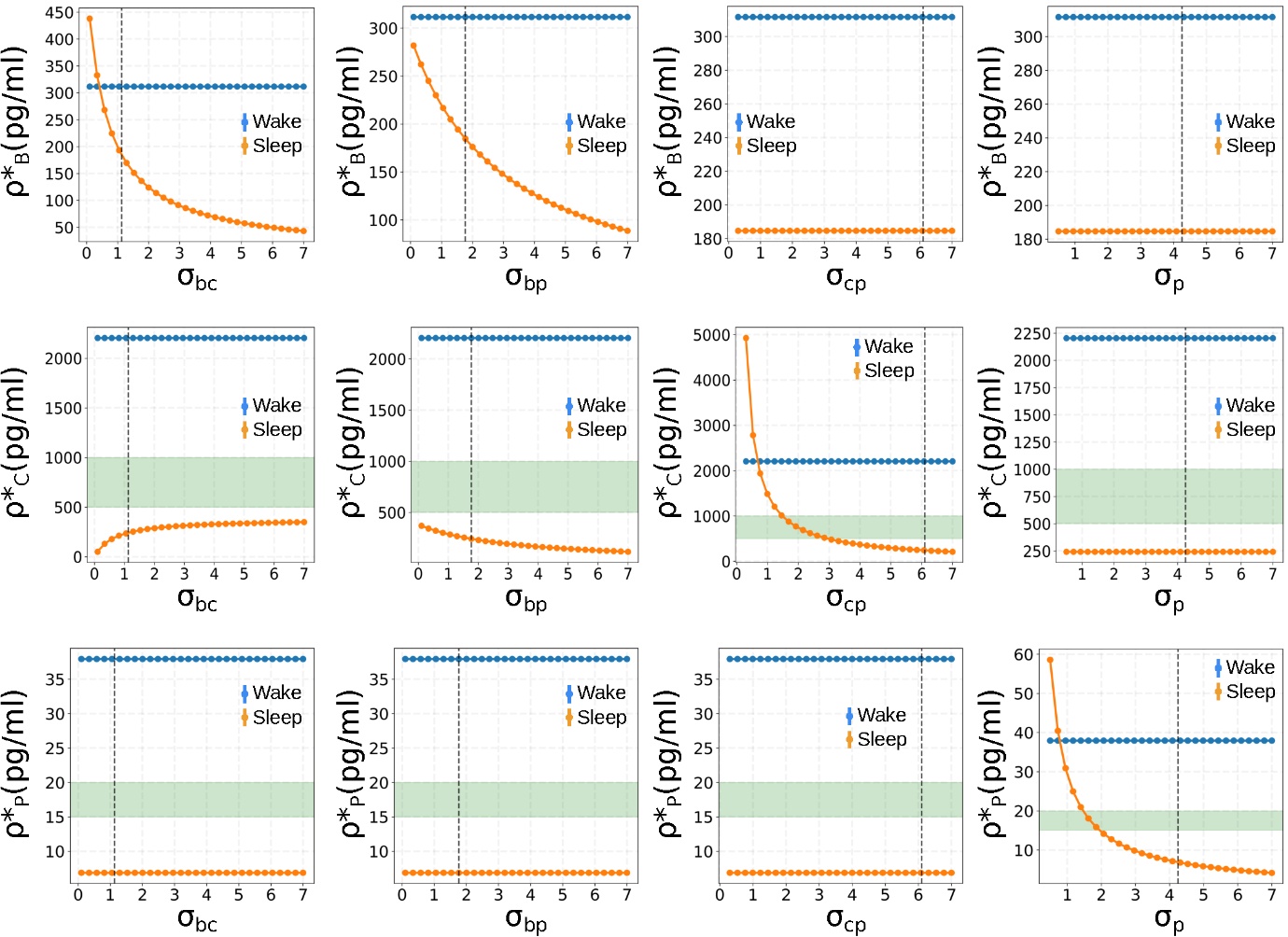

**Figure S2: Steady state response of the respective compartments during wake (blue curve) and sleep (orange curve) state**. (top row) Brain compartment vs sleep state parameters $\sigma_{bc}$, $\sigma_{bp}$*,* $\sigma_{cp}$*,* $\sigma_{p}$ (middle-row) CSF compartment vs sleep state parameters $\sigma_{bc}$, $\sigma_{bp}$*,* $\sigma_{cp}$*,* $\sigma_{p}$ (bottom-row) Plasma compartment vs sleep state parameters $\sigma_{bc}$, $\sigma_{bp}$*,* $\sigma_{cp}$*,* $\sigma_{p}$

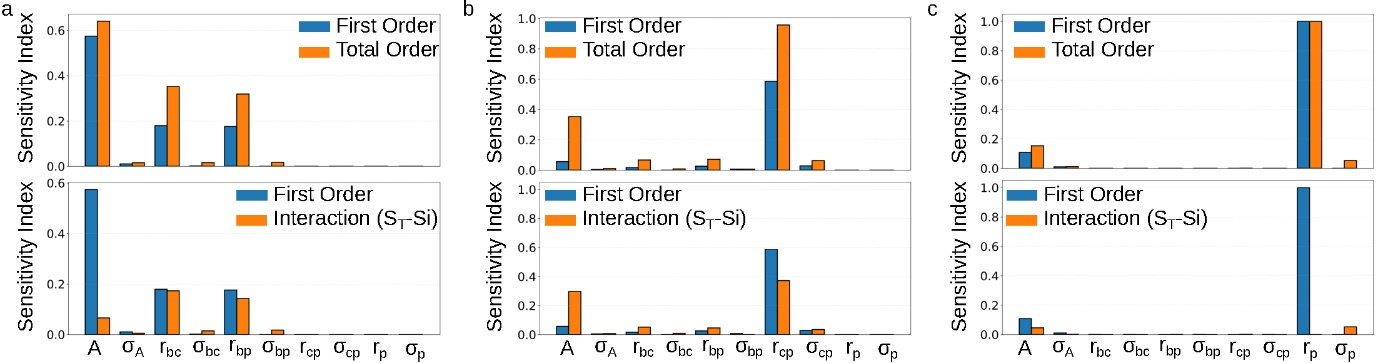

**Figure S3:** **Global sensitivity analysis of the Aβ42 compartmental model using Sobol indices.** The upper panel shows first-order ($S_{i}$, blue) and total-order ($S_{T}$, orange) Sobol sensitivity indices for model outputs in the brain (a), CSF (b), and plasma (c) compartments. The lower panel shows the interaction contribution ($S_{T}-S_{i}$, orange) for each parameter, representing the proportion of output variance attributable to parameter interactions.

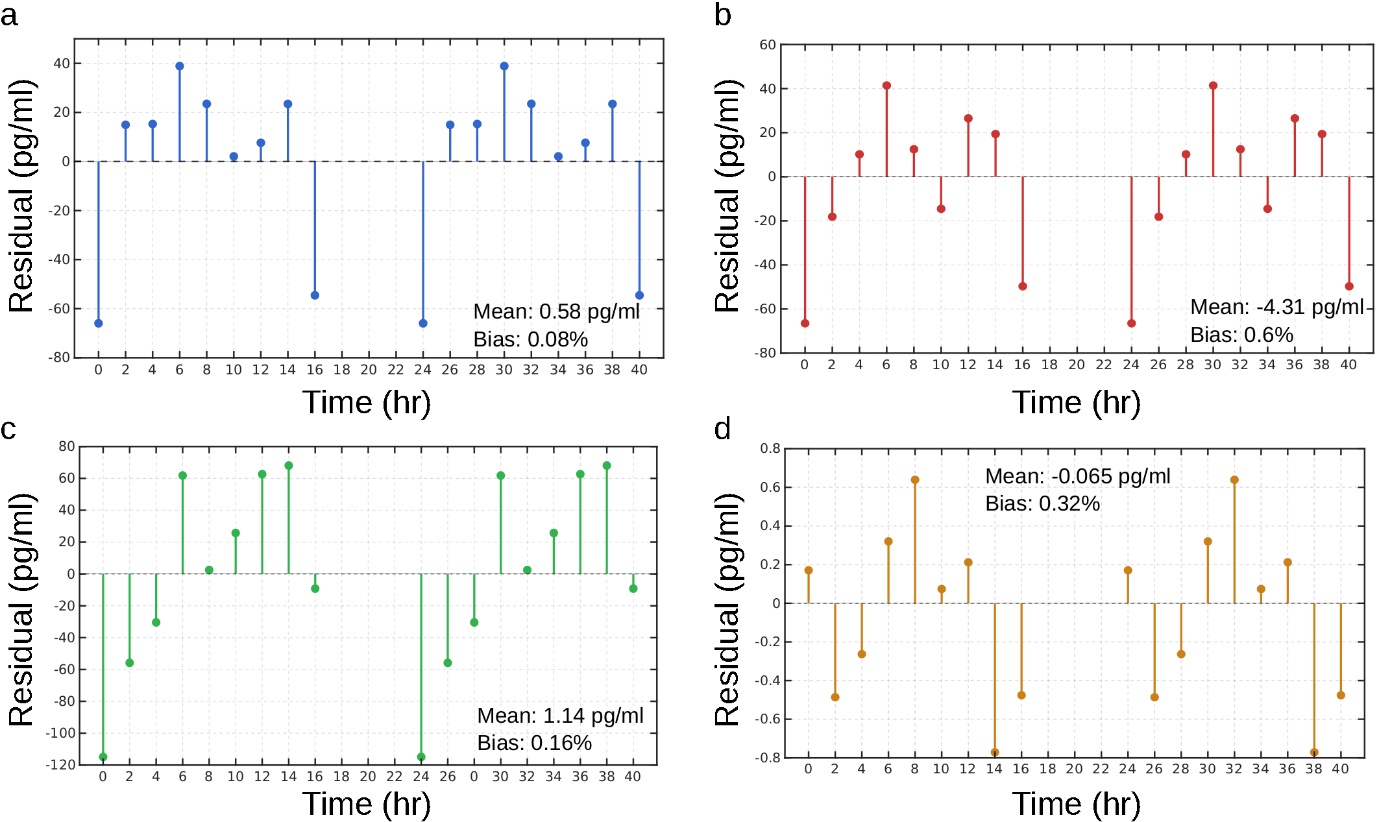

**Figure S4:** **Residual analysis of the fitted Aβ42 model.** Residuals are shown for (a) Blattner et al. 2020 (b) Lucey et al. 2018 (c) Liu et al. 2022 CSF data and (d) Liu et al. 2022 plasma data along with their mean and bias.

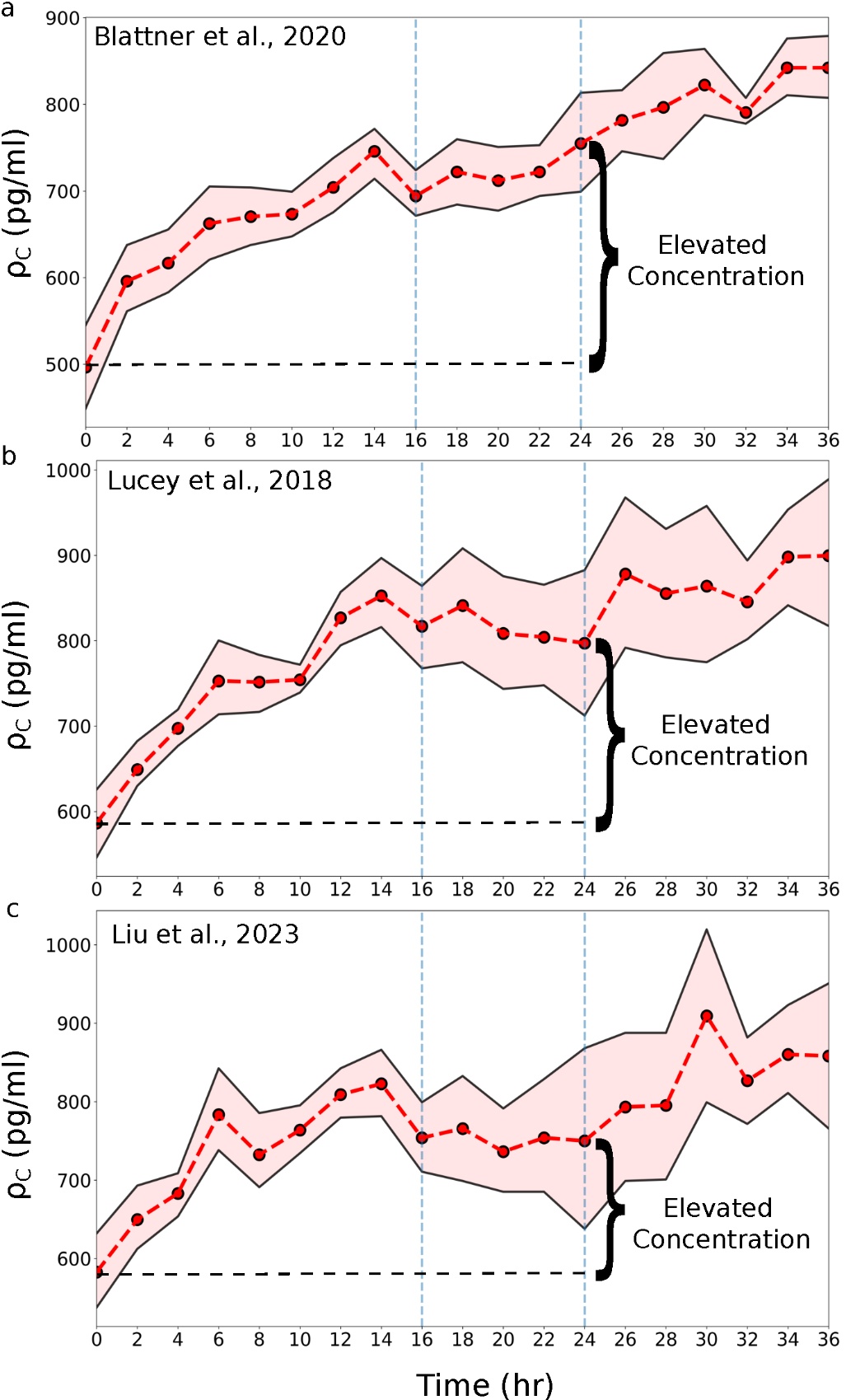

**Figure S5: CSF Amyloid levels for the experimental dataset showing the sampling discrepancy in CSF amyloid levels** (a) Blattner et al.^1^ (b) Lucey et al.^2^ (c) Liu et al.^3^

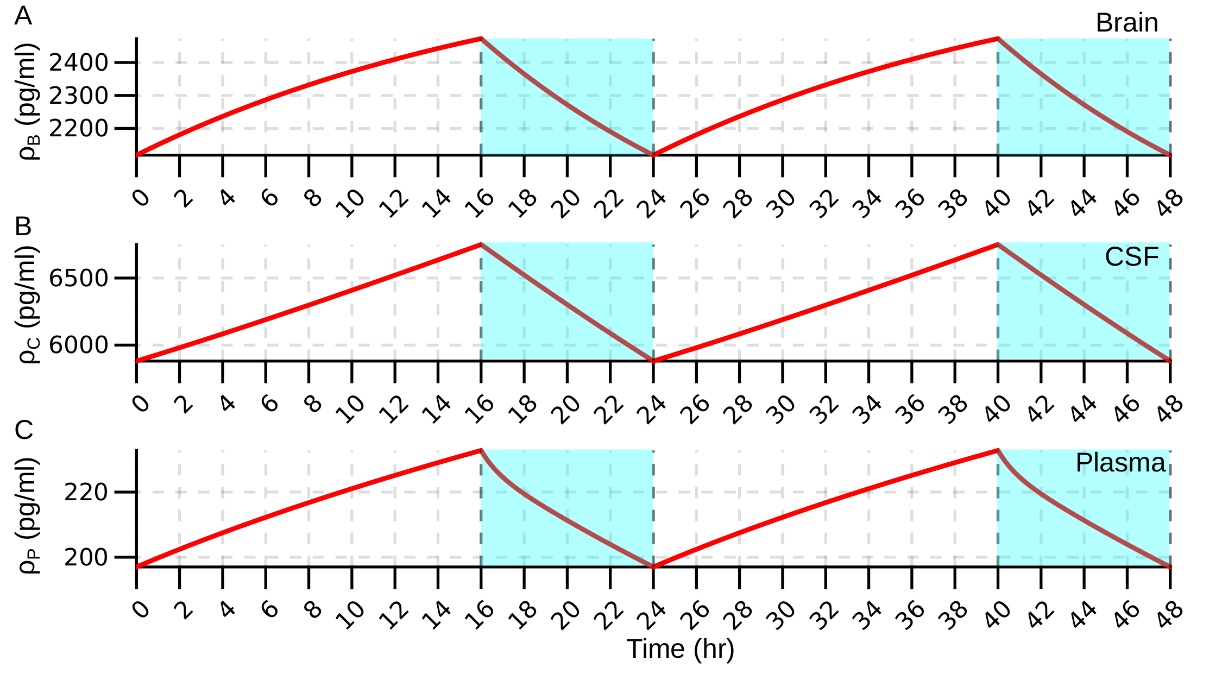

**Figure S6: Model Aβ40 Dynamics over a Period of 48 Hours**. **(A)** The brain amyloid concentration fluctuates between 2542-2040 pg/ml. **(B)** The CSF amyloid concentration fluctuates between 6987-5956 pg/ml. **(C)** The plasma amyloid concentration fluctuates between 234-202 pg/ml. The cyan shaded area corresponds to sleep state. The vertical dashed line corresponds to the wake-to-sleep transition (t=16 hr, t=40 hr) and the sleep-to-wake transition (t=24 hr, t=48 hr)

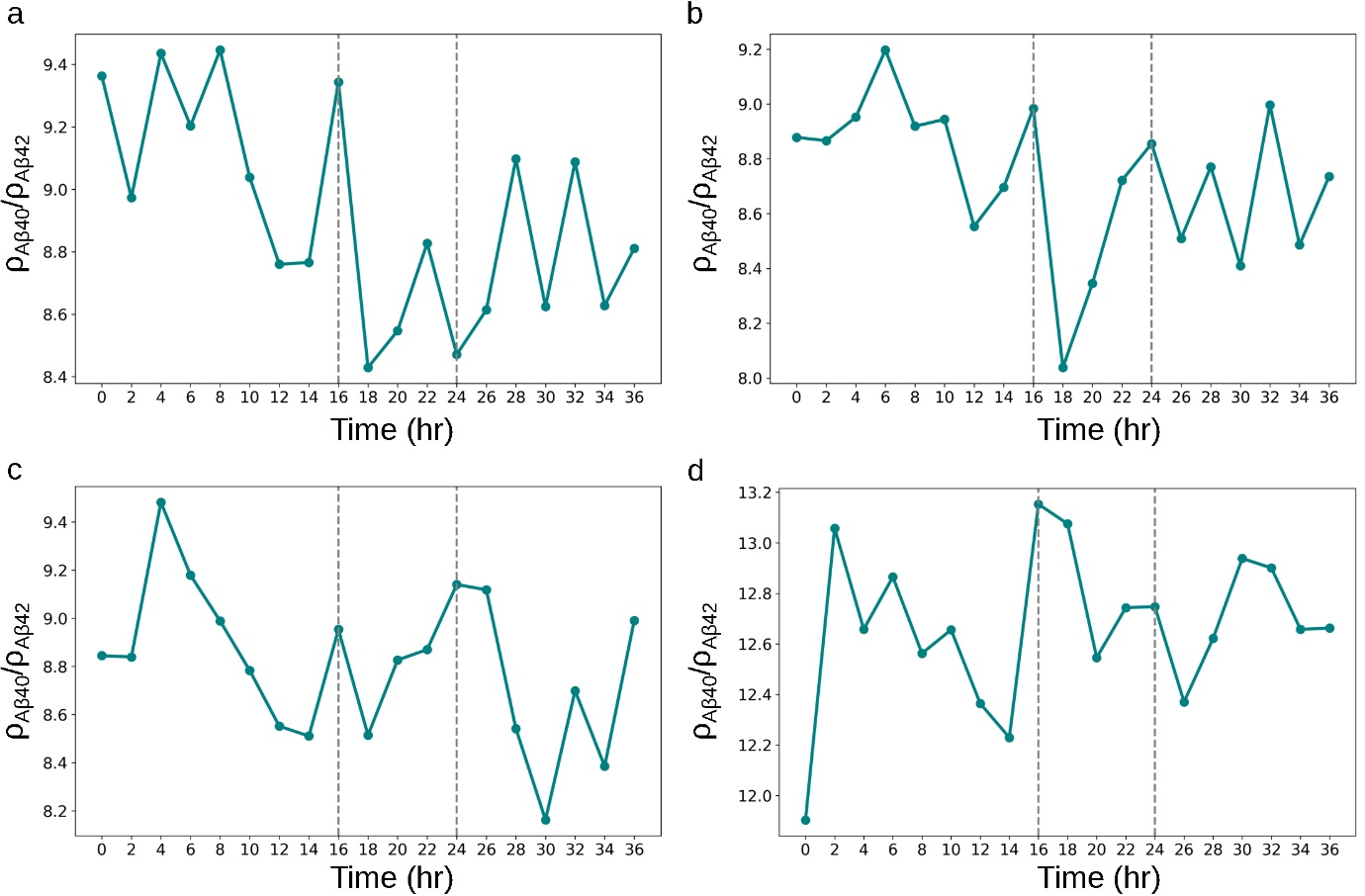

**Figure S7: Ratio of concentration Aβ40 to Aβ42 in CSF and plasma compartments.** The line plot shows the temporal concentration evolution of ratio of Aβ40:Aβ42 in (a) Blattner et al.^1^ (b) Lucey et al.^2^ (c) CSF compartment of Liu et al.^3^ and (d) Plasma compartment of Liu et al.^3^

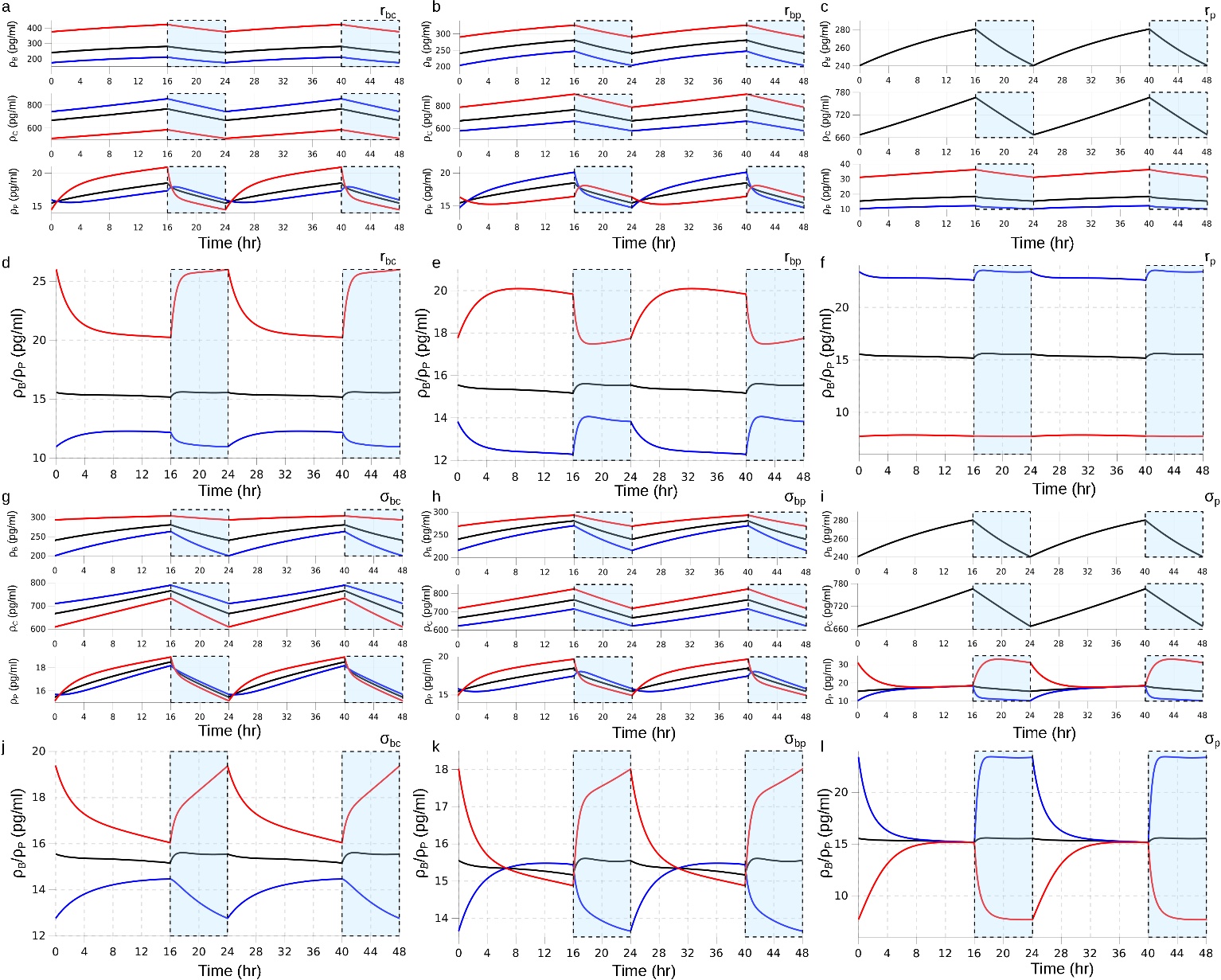

**Figure S8: Model predicted brain-plasma Aβ relationship** showing 48-hour model trajectories of brain, CSF and plasma Aβ concentrations under systematic variation of clearance rate constants (a) $r_{bc}$ (b) $r_{bp}$ (c) $r_{p}$ and sleep-dependent scaling factors (g) $\sigma_{bc}$ (h) $\sigma_{bp}$ (i) $\sigma_{p}$. Each panel shows three representative parameter values with parameters at their default value (black), 50% decreased (red), and 50% increased (blue) with all other parameters held at their globally fitted values. Panels d–f and j–l shows the corresponding brain-to-plasma ratio (${\rho_{b}}/{\rho_{p}}$) for each parameter variation. Vertical dashed lines represent sleep onset (hour 16, hour 40) and wake onset (hour 24, hour 48).

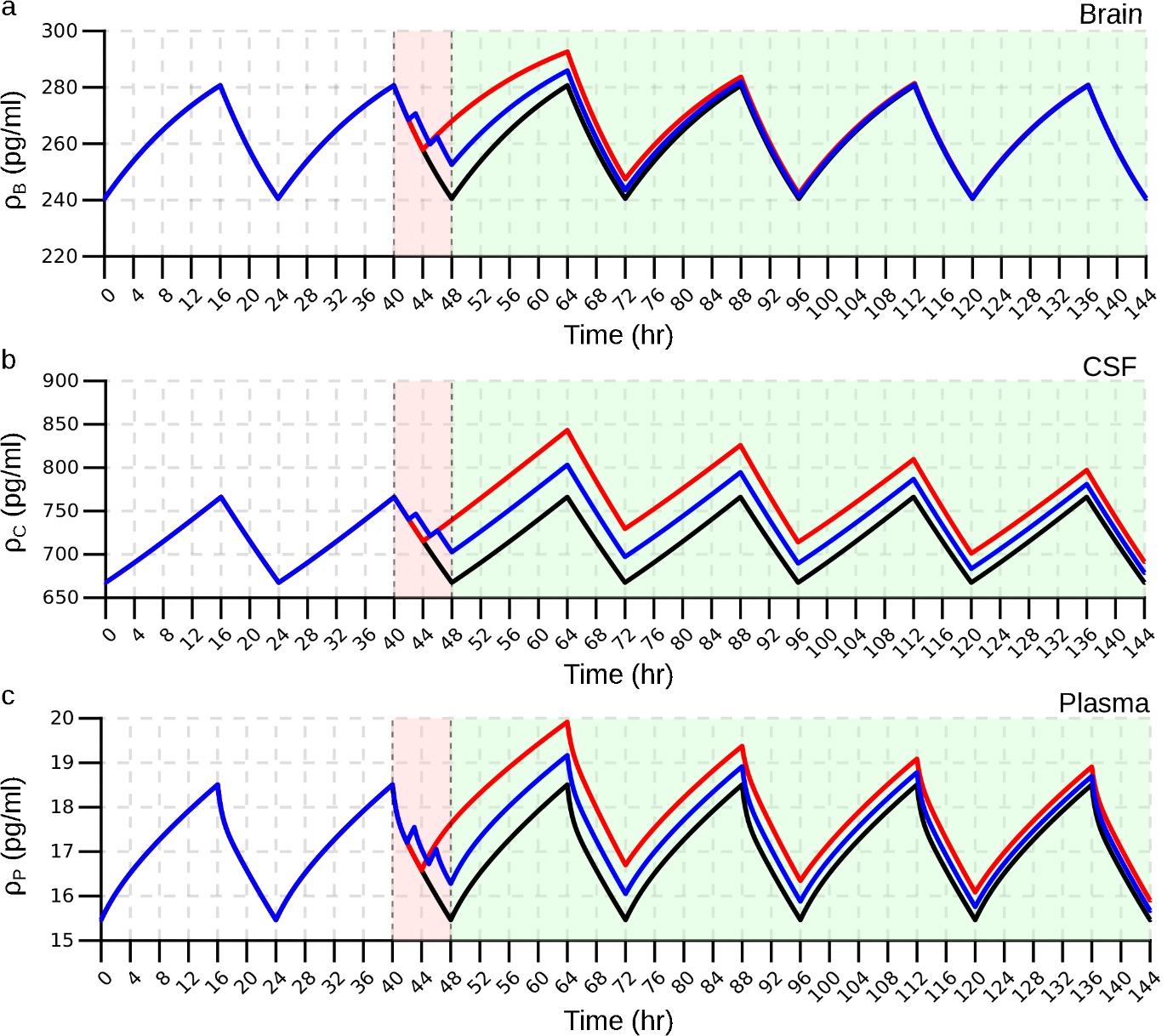

**Figure S9: Model prediction of Aβ dynamics** in (a) brain (b) CSF and (c) plasma compartment for normal (8 hours sleep, black), partial (4 hours sleep, red) and fragmented (three 2 hours sleep cycle totalling 6 hours with two 1 hour of wake cycle in between, blue) sleep. Pink shaded region indicates the intervention sleep window during which the three protocols are applied. Green shaded region represents the post-intervention period when model falls back to normal sleep-wake cycle.

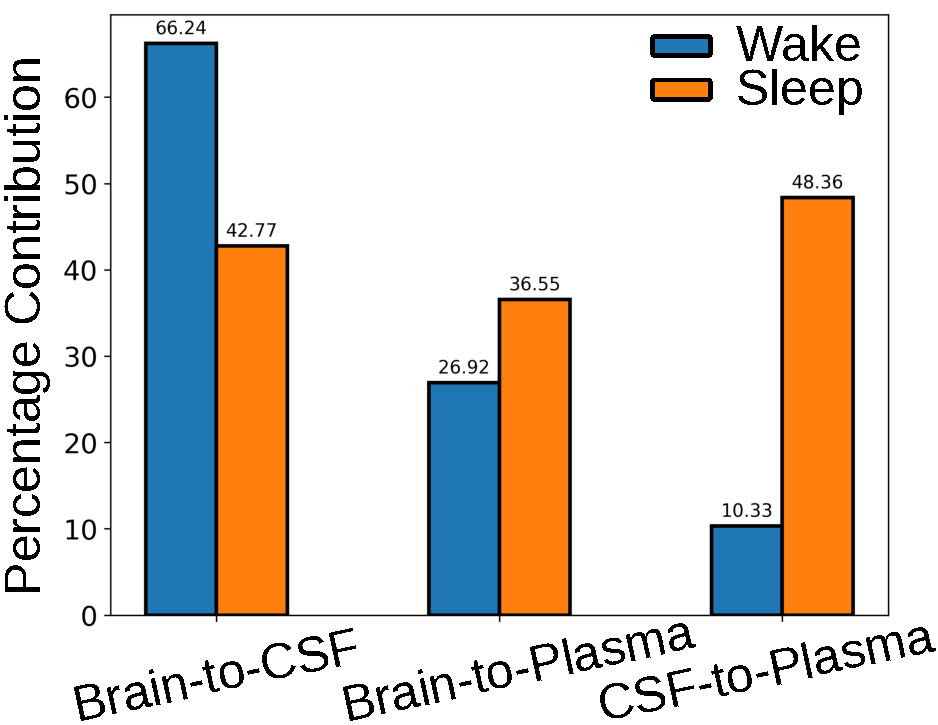

**Figure S10: Contribution of individual pathways to the total CNS clearance.** The bars correspond to the total flux contribution ($r_{i}\sigma_{i}\rho_{i}$) by individual pathways during wake (blue) and sleep (orange) state.

**Table S1: Changes in NRMSE and AICc for lumbar hypothesis testing for Aβ40.** Hypothesis NRMSE (percentage of baseline, %) represents the percentage change in fit quality compared to baseline NRMSE. For the global column, baseline is the global fit across all datasets using default parameters. For individual dataset columns (Blattner et al.^1^, Lucey et al.^2^, Liu et al.^3^), baseline is the respective individual fits from normal sleep-wake fitting. Negative values indicate the hypothesis improved the fit quality compared to baseline, positive values indicate worsened fit and zero indicates no change. The lowest values for each dataset are highlighted in bold.

| Parameter Combination | Hypothesis NRMSE change compared to baseline NRMSE (%) | | | | Corrected AIC | | | |
| --- | --- | --- | --- | --- | --- | --- | --- | --- |
|  | **Global** | **Blattner et al.^1^** | **Lucey et al.^2^** | **Liu et al.^3^** | **Global** | **Blattner et al.^1^** | **Lucey et al.^2^** | **Liu et al.^3^** |
| Pressure Hypothesis | | | | | | | | |
| $\boldsymbol{r}_{\boldsymbol{bc}}$ | -22 | -44 | -25 | -9 | 1041.42 | 299.10 | 300.73 | 440.65 |
| $\boldsymbol{r}_{\boldsymbol{cp}}$ | -1 | -63 | -43 | 0 | 1059.38 | 283.95 | 290.19 | 447.59 |
| $\boldsymbol{r}_{\boldsymbol{bc}}\boldsymbol{,}\boldsymbol{r}_{\boldsymbol{cp}}$ | -22 | -63 | -44 | -9 | 1043.78 | 286.86 | 292.96 | 443.16 |
| $\boldsymbol{r}_{\boldsymbol{bp}}\boldsymbol{,}\boldsymbol{r}_{\boldsymbol{p}}$ | -2 | -58 | -37 | -10 | 1050.48 | 291.16 | 297.26 | 441.30 |
| $\boldsymbol{r}_{\boldsymbol{bc}}\boldsymbol{,}\boldsymbol{r}_{\boldsymbol{cp}}\boldsymbol{,}\boldsymbol{r}_{\boldsymbol{bp}}$ | -27 | **-70** | **-59** | -21 | 1028.24 | **281.50** | **284.44** | 435.48 |
| $\boldsymbol{r}_{\boldsymbol{bc}}\boldsymbol{,}\boldsymbol{r}_{\boldsymbol{cp}}\boldsymbol{,}\boldsymbol{r}_{\boldsymbol{p}}$ | **-37** | -62 | -43 | **-30** | **1013.36** | 290.21 | 296.58 | **425.65** |
| Sleep Hypothesis | | | | | | | | |
| $\boldsymbol{\sigma}_{\boldsymbol{bc}}$ | 0 | 0 | 0 | 0 | 1059.38 | 321.34 | 311.84 | 447.59 |
| $\boldsymbol{\sigma}_{\boldsymbol{cp}}$ | 0 | -65 | -53 | 0 | 1059.38 | **280.90** | **282.96** | 447.59 |
| $\boldsymbol{\sigma}_{\boldsymbol{A}}$ | -3 | -13 | -30 | 0 | 1058.78 | 315.86 | 298.20 | 447.20 |
| $\boldsymbol{\sigma}_{\boldsymbol{bp}}$ | 0 | -41 | -45 | 0 | 1059.38 | 301.09 | 288.96 | 447.59 |
| $\boldsymbol{\sigma}_{\boldsymbol{p}}$ | -3 | 0 | 0 | -12 | 1047.34 | 321.32 | 311.84 | 435.55 |
| $\boldsymbol{\sigma}_{\boldsymbol{A}}\boldsymbol{,}\boldsymbol{\sigma}_{\boldsymbol{p}}$ | -3 | -13 | -30 | -12 | 1049.70 | 318.71 | 301.05 | 438.06 |
| $\boldsymbol{\sigma}_{\boldsymbol{bc}}\boldsymbol{,}\boldsymbol{\sigma}_{\boldsymbol{cp}}$ | 0 | **-68** | **-54** | 0 | 1061.74 | 281.24 | 285.12 | 450.09 |
| $\boldsymbol{\sigma}_{\boldsymbol{A}}\boldsymbol{,}\boldsymbol{\sigma}_{\boldsymbol{bp}}$ | -3 | -54 | -46 | 0 | 1061.16 | 294.45 | 291.14 | 449.72 |
| $\boldsymbol{\sigma}_{\boldsymbol{bp}}\boldsymbol{,}\boldsymbol{\sigma}_{\boldsymbol{p}}$ | -3 | -41 | -45 | -12 | 1049.70 | 303.94 | 291.81 | 438.06 |
| $\boldsymbol{\sigma}_{\boldsymbol{cp}}\boldsymbol{,}\boldsymbol{\sigma}_{\boldsymbol{p}}$ | **-32** | -35 | -53 | **-26** | **1017.67** | 283.75 | 285.81 | **428.04** |
| $\boldsymbol{\sigma}_{\boldsymbol{bc}}\boldsymbol{,}\boldsymbol{\sigma}_{\boldsymbol{bp}}\boldsymbol{,}\boldsymbol{\sigma}_{\boldsymbol{cp}}$ | 0 | **-68** | **-54** | 0 | 1064.17 | 284.49 | 288.38 | 452.76 |
| $\boldsymbol{\sigma}_{\boldsymbol{A}}\boldsymbol{,}\boldsymbol{\sigma}_{\boldsymbol{bc}}\boldsymbol{,}\boldsymbol{\sigma}_{\boldsymbol{cp}}$ | -3 | **-68** | **-54** | 0 | 1063.59 | 284.49 | 288.38 | 452.38 |
| $\boldsymbol{\sigma}_{\boldsymbol{bc}}\boldsymbol{,}\boldsymbol{\sigma}_{\boldsymbol{cp}}\boldsymbol{,}\boldsymbol{\sigma}_{\boldsymbol{p}}$ | **-32** | **-68** | **-54** | **-26** | 1019.91 | 284.49 | 288.38 | 430.65 |
| $\boldsymbol{\sigma}_{\boldsymbol{A}}\boldsymbol{,}\boldsymbol{\sigma}_{\boldsymbol{bp}}\boldsymbol{,}\boldsymbol{\sigma}_{\boldsymbol{p}}$ | -3 | -54 | -46 | -12 | 1052.13 | 297.71 | 294.40 | 440.72 |
| $\boldsymbol{\sigma}_{\boldsymbol{A}}\boldsymbol{,}\boldsymbol{\sigma}_{\boldsymbol{bc}}\boldsymbol{,}\boldsymbol{\sigma}_{\boldsymbol{cp}}\boldsymbol{,}\boldsymbol{\sigma}_{\boldsymbol{p}}$ | **-32** | **-68** | **-54** | **-26** | 1022.41 | 288.25 | 292.14 | 433.49 |
| $\boldsymbol{\sigma}_{\boldsymbol{A}}\boldsymbol{,}\boldsymbol{\sigma}_{\boldsymbol{bc}}\boldsymbol{,}\boldsymbol{\sigma}_{\boldsymbol{bp}}\boldsymbol{,}\boldsymbol{\sigma}_{\boldsymbol{cp}}$ | -3 | **-68** | **-54** | 0 | 1066.09 | 288.25 | 292.14 | 455.21 |
| $\boldsymbol{\sigma}_{\boldsymbol{bc}}\boldsymbol{,}\boldsymbol{\sigma}_{\boldsymbol{bp}}\boldsymbol{,}\boldsymbol{\sigma}_{\boldsymbol{cp}}\boldsymbol{,}\boldsymbol{\sigma}_{\boldsymbol{p}}$ | **-32** | **-68** | **-54** | **-26** | 1022.41 | 288.25 | 292.14 | 433.49 |
| $\boldsymbol{\sigma}_{\boldsymbol{A}}\boldsymbol{,}\boldsymbol{\sigma}_{\boldsymbol{bc}}\boldsymbol{,}\boldsymbol{\sigma}_{\boldsymbol{bp}}\boldsymbol{,}\boldsymbol{\sigma}_{\boldsymbol{cp}}\boldsymbol{,}\boldsymbol{\sigma}_{\boldsymbol{p}}$ | **-32** | **-68** | **-54** | **-26** | 1024.99 | 292.63 | 296.53 | 436.51 |
| Combined Hypothesis | | | | | | | | |
| $\boldsymbol{r}_{\boldsymbol{bc}}\boldsymbol{,}\boldsymbol{\sigma}_{\boldsymbol{bc}}\boldsymbol{,}\boldsymbol{\sigma}_{\boldsymbol{cp}}$ | -22 | -68 | -54 | -9 | 1046.23 | **284.48** | **288.38** | 445.83 |
| $\boldsymbol{r}_{\boldsymbol{cp}}\boldsymbol{,}\boldsymbol{\sigma}_{\boldsymbol{bc}}\boldsymbol{,}\boldsymbol{\sigma}_{\boldsymbol{cp}}$ | 0 | -68 | -54 | 0 | 1064.17 | **284.48** | **288.38** | 452.76 |
| $\boldsymbol{r}_{\boldsymbol{bc}}\boldsymbol{,}\boldsymbol{\sigma}_{\boldsymbol{cp}}\boldsymbol{,}\boldsymbol{\sigma}_{\boldsymbol{p}}$ | **-39** | -65 | -53 | **-33** | **1004.38** | 287.01 | 289.07 | **422.50** |
| $\boldsymbol{r}_{\boldsymbol{bc}}\boldsymbol{,}\boldsymbol{r}_{\boldsymbol{cp}}\boldsymbol{,}\boldsymbol{\sigma}_{\boldsymbol{bc}}\boldsymbol{,}\boldsymbol{\sigma}_{\boldsymbol{cp}}$ | -22 | -68 | -54 | -9 | 1048.73 | 288.24 | 292.14 | 448.66 |
| $\boldsymbol{r}_{\boldsymbol{bc}}\boldsymbol{,}\boldsymbol{\sigma}_{\boldsymbol{bc}}\boldsymbol{,}\boldsymbol{\sigma}_{\boldsymbol{cp}}\boldsymbol{,}\boldsymbol{\sigma}_{\boldsymbol{p}}$ | **-39** | -68 | -54 | **-33** | 1006.91 | 288.25 | 292.14 | 425.34 |
| ${\boldsymbol{r}_{\boldsymbol{bc}}\boldsymbol{,}\boldsymbol{r}_{\boldsymbol{bp}}\boldsymbol{,r}}_{\boldsymbol{cp}}\boldsymbol{,}$  $\boldsymbol{\sigma}_{\boldsymbol{bc}}\boldsymbol{,}\boldsymbol{\sigma}_{\boldsymbol{bp}}\boldsymbol{,}\boldsymbol{\sigma}_{\boldsymbol{cp}}$ | -29 | **-76** | **-63** | -20 | 1043.05 | 286.45 | 293.53 | 445.50 |
| ${\boldsymbol{r}_{\boldsymbol{bc}}\boldsymbol{,}\boldsymbol{r}_{\boldsymbol{cp}}\boldsymbol{,}\boldsymbol{\sigma}_{\boldsymbol{A}}\boldsymbol{,\sigma}}_{\boldsymbol{bc}}\boldsymbol{,}$  $\boldsymbol{\sigma}_{\boldsymbol{bp}}\boldsymbol{,}\boldsymbol{\sigma}_{\boldsymbol{cp}}\boldsymbol{,}\boldsymbol{\sigma}_{\boldsymbol{p}}$ | **-39** | -68 | -54 | **-33** | 1006.91 | 304.02 | 307.93 | 435.06 |

**Table S2: Profile likelihood analysis for the model parameters showing the optimum values and the confidence intervals**

| Parameter | Optimum Value | Lower Bound | Upper Bound | Interval |
| --- | --- | --- | --- | --- |
| $\boldsymbol{r}_{\boldsymbol{bc}}$ | 0.038 | 0.01 | 0.25 | [0.026,0.055] |
| $\boldsymbol{r}_{\boldsymbol{bp}}$ | 0.014 | 0.01 | 0.25 | [0.01,0.019] |
| $\boldsymbol{r}_{\boldsymbol{cp}}$ | 0.00537 | 0 | 0.1 | [0.00497,0.00597] |
| $\boldsymbol{\sigma}_{\boldsymbol{bc}}$ | 1.131 | 1 | 7 | [1,2.68] |
| $\boldsymbol{\sigma}_{\boldsymbol{bp}}$ | 1.768 | 1 | 7 | [1,3.080] |
| $\boldsymbol{\sigma}_{\boldsymbol{cp}}$ | 6.100 | 1 | 7 | [5.311,7] |
| $\boldsymbol{\sigma}_{\boldsymbol{p}}$ | 4.253 | 1 | 7 | [3.291,6.065] |
| $\boldsymbol{A}$ | 16.203 | 0 | 111 | [15.06,17.291] |
| $\boldsymbol{\sigma}_{\boldsymbol{A}}$ | 0.772 | 0 | 0.99 | [0.537,0.980] |
| $\boldsymbol{r}_{\boldsymbol{p}}$ | 0.427 | 0 | 0.6 | [0.391,0.479] |

**Table S3: Experimental studies used for calculating Amyloid concentration in CSF and Plasma for cognitively healthy participants.** The data is obtained from Mo et al., 2015^4^

| Study | Age (years) | Mean ± SD (pg/ml) | Assay Type |
| --- | --- | --- | --- |
| CSF Data |  |  |  |
| Kapaki et al.^5^ (N=50) | 62±12* | 736±157 | Double sandwich ELISA using ‘Innotest htau antigen’ and ‘Innotest beta-amyloid (1-42)’ kits. Innogenetics |
| Kapaki et al.^6^ (N=64) | 64±11* | 721±228 | Double sandwich ELISA using ‘Innotest htau antigen’ and ‘Innotest beta-amyloid (1-42)’ kits. Innogenetics |
| Reijin et al.^7^ (N=55) | 59 (53-67)^#^ | 810±170 (E)  381±68(X) | CSF were measured with both ELISA (E) and xMAP-based (X) Innobia assay (Innogenetics) |
| Lewczuk et al.^8^ (N=35) | 61 (52-67)^#^ | 865±256 | ELISA test kits |
| Schoonenboom et al.^9^ (N=21) | 62 (39-70)^$^ | 604±443.5 | Sandwich ELISA (Innotest beta-amyloid) |
| Le Bastard et al.^10^ (N=95) | 47±17* | 699±417(IT)  212±116.5(IB) | Amyloid levels in CSF were measured with commercially available single analyte ELISA kits (Innotest, IT) and with a research use only multi-analyte Luminex assay (Inno-Bia, IB) |
| Buchhave et al.^11^ (N=34) | 72±8.3* | 651±178 | Sandwich ELISA (Innotest) was used for AD Cohorts and xMAP (INNO-BIA) was used for Control and then the results were converted to ELISA levels using the conversion factor* |
| Smach et al.^12^ (N=35) | 72 (55-87)^ʈ^ | 1020±230 | ELISA (Innotest) |
| Mattsson et al.^13^ (N=304) | 67 (44-91)^ʈ^ | 675±285.8 | Sandwich ELISA (Innotest). For two medical centers (Malmo and Goteberg) xMAP was used and the data was converted to ELISA level |
| Andreasen et al.^14^ (N=21) | 68.8±8* | 1678±436 | Sandwich ELISA |
| Plasma Data |  |  |  |
| Zecca et al.^15^ (N=93) | 75.40±6.78*  (>65 years) | 18.67±6.36 | ELISA (Innotest) |
| Feinkohl et al.^16^ (N=50) | 65.82±8.96* | 19.9±6.6 | ELISA Assay |
| Ruiz et al.^17^ (N=53) | 60.3±8.3* | 13±12 (DA)  46.5±28 (RP)  149.1±69 (CP) | Sandwich ELISA |
| Pesini et al.^18^ (N=16) | 70.3±4.1* | 8±3.09 (UP)  40.1±12.91 (DP)  58.7±13.6 (CB) | Sandwich ELISA |

**Mean±SD; ^#^Median value (p25-p75); ^$^Median value (Minimum-Maximum); ^ʈ^Median (Range); ^†^CSF amyloid Data of these studies are originally presented as median and percentile ranges (p25-p75); ^‡^CSF amyloid Data of these studies are originally presented as medians (minimum-maximum); ^⁑^CSF amyloid Data of these studies are originally presented as Mean (Coefficient of Variation). All of these were converted to SD (Details about conversion is provided in supplementary)*

***Estimation of Concentration Ranges and Averages***

*Inter-Quartile Range (IQR) Method*

To determine the theoretical range of Aβ concentration in plasma and CSF, we computed the interquartile range (IQR) using distributions of experimental data^19^. We applied a multiplier of 2 instead of 1.5 to broaden the range and accommodate wider physiological fluctuations in Aβ levels.

$$Range=\left[ Q_{1}-1.5*IQR, Q_{3}+1.5*IQR \right] eq(S1)$$

***CSF Aβ42 Range Calculation:***

Ordered Mean data: [604, 659, 675, 699, 721, 736, 810, 865, 1020] $pg{ml}^{-1}$ (n=9)

Q1 (25^th^ percentile): 667 $pg{ml}^{-1}$

Q3 (75^th^ percentile): 837.5 $pg{ml}^{-1}$

IQR = 170.5 $pg{ml}^{-1}$, Range = [326, 1179] ≈ [320, 1200] $pg{ml}^{-1}$

***Plasma Aβ42 Range Calculation:***

Ordered Mean data: [8, 13, 18.67, 19.9] $pg{ml}^{-1}$ (n=4)

Q1 (25^th^ percentile): 9.25 $pg{ml}^{-1}$

Q3 (75^th^ percentile): 19.59 $pg{ml}^{-1}$

IQR = 10.34 $pg{ml}^{-1}$, Range = [0, 41]

***Decomposition of Mean and Standard Deviation (DMSD) Method***

We used the DMSD^20^ method to aggregate data across multiple studies with varying sample sizes, mean, and standard deviations. For each study:

$$\sum x=\mu.n eq(S2)$$

$$\sum x^{2}=\sigma^{2}.\left( n-1 \right)+\frac{{(\sum x)}^{2}}{n} eq(S3)$$

Combined estimates were calculated from aggregated values ($t_{x},t_{xx},t_{n}$):

$$Combined Mean=\frac{t_{x}}{t_{n}}, Combined SD=\sqrt{\frac{t_{xx}-\frac{{(t_{x})}^{2}}{t_{n}}}{t_{n}-1}} eq(S4)$$

**Final combined estimates:**

CSF Aβ42 = 721 ± 294.26 $pg{ml}^{-1}$

CSF Aβ40 = 6404.19 ± 4548.96 $pg{ml}^{-1}$

Plasma Aβ42 = 16.74 ± 8.79 $pg{ml}^{-1}$

Plasma Aβ42 = 213.95 ± 91.96 $pg{ml}^{-1}$

**Standard Deviation Conversions**

Studies reported variability using different statistical measures. We applied the following conversions to standardize all data to mean ± SD format:

*From Standard Error:*

$$\sigma=SE.\sqrt{N} eq(S5)$$

*From Interquartile Range:*

$$\sigma=\frac{{IQR}_{75}-{IQR}_{25}}{2} eq(S6)$$

*From Range:*

For N ≤ 30: $\sigma\approx(Max-Min)/4$

For N $>$ 30: $\sigma\approx(Max-Min)/6$

*From Coefficient of Variation:*

$$\sigma=\frac{CV.\mu}{100} eq(S7)$$

These conversions assume normally distributed data and may be less accurate for significantly skewed distributions.

### **Sensitivity Analysis**

Local sensitivity analysis was performed to quantify the effect of small perturbations in individual model parameters on the predicted Aβ concentrations in the brain, CSF and plasma compartments. The model was first simulated using globally optimised parameters set to establish baseline Aβ concentration profile over a 24-hour period. Each parameter was then independently increased by 1% while all other parameters were held constant, and the model was re-simulated. The sensitivity of each compartmental output to each parameter was calculated over the 24 hours period by summing the compartment concentrations within this 24-hour window for both the baseline and perturbed simulations. The normalised local sensitivity coefficient for output $y_{i}$ (summed Aβ concentration in a compartment) with respect to parameter $p_{j}$ was calculated as:

$$S_{i,j}=\frac{\frac{\Delta y_{i}}{y_{i}}}{\frac{\Delta p_{j}}{p_{j}}} eq(S8)$$

Where $\Delta y_{i}$ is the change in output resulting from a small change $\Delta p_{j}$ in parameter $p_{j}$, where $y_{i}$ and $p_{j}$ are the respective baseline values. This dimensionless coefficient represents the percentage change in output per percentage change in the parameter, allowing a direct comparison across parameters with different units and magnitudes.
